# Rab7/Retromer-based endolysosomal trafficking facilitates effector secretion and host invasion in rice blast

**DOI:** 10.1101/2022.06.22.497169

**Authors:** Xin Chen, Poonguzhali Selvaraj, Lili Lin, Wenqin Fang, Congxian Wu, Piao Yang, Jing Zhang, Yakubu Saddeeq Abubakar, Fan Yang, Guodong Lu, Wende Liu, Zonghua Wang, Naweed I. Naqvi, Wenhui Zheng

**Affiliations:** State Key Laboratory for Ecological Pest Control of Fujian and Taiwan Crops, College of Plant Protection, Fujian Agriculture and Forestry University, Fuzhou, China; Key Laboratory of Bio-pesticide and Chemistry Biology, Ministry of Education, College of Plant Protection, Fujian Agriculture and Forestry University, Fuzhou, Fujian, China; Temasek Life Sciences Laboratory, and the Department of Biological Sciences, National University of Singapore, Singapore, Singapore; State Key Laboratory for Biology of Plant Diseases and Insect Pests, Institute of Plant Protection, Chinese Academy of Agricultural Sciences, Beijing, China; Institute of Oceanography, Minjiang University, Fuzhou, China

## Abstract

Secretion is a fundamental process in all living organisms. Using conventional secretion pathways, many plant pathogens release effectors into the host plants to downregulate immunity and promote infection. However, this does not always constitute the only way that effectors are sorted and tracked to their final destination such as the biotrophic interfacial complex-associated effectors produced by the blast fungus *Magnaporthe oryzae.* Here, we uncover a novel unconventional route originating from fungal vacuolar membrane to the host interface and plasma membrane. We found that a GFP-MoRab7 labeled vacuole is closely associated with the interface structure throughout *M. oryzae* invasive growth. Conditional inactivation of MoRab7 impaired the establishment of the biotrophy interface and secretion of Pwl2 effector. To perform the vacuolar trafficking pathway, MoRab7 first recruits the retromer complex to the vacuole membrane, enabling it recognizes a batch of SNARE proteins, including the v-SNARE MoSnc1. Live-cell imaging supports both retromer complex component and MoSnc1 protein labeled vesicles showing the trafficking dynamics toward the interface or plasma membrane, and then fusion with target membranes. Lastly, disruption of the MoRab7/Retromer/MoSnc1-based endolysosomal cascade affects effector secretion and fungal pathogenicity. Taken together, we discovered an unconventional protein and membrane trafficking route starting from the fungal endolysosomes to the *M. oryzae*-rice interaction interface, and dissect the role of MoRab7/Retromer/MoSnc1 constituent sorting machinery in effector secretion during invasive growth in *M. oryzae*.

## Introduction

Each year, crop diseases caused by phytopathogenic fungi lead to extensive yield loss and seriously threaten global food security [1, 2]. Despite the differences in lifestyles, a majority of these phytopathogens use a common strategy i.e. secretion of fungal effectors into the host to facilitate colonization [2, 3]. *M. oryzae*, the causal agent of the rice blast disease, seriously threatens rice production worldwide and has emerged as a top model to study host-fungus interaction [1, 4]. During host interaction, *M. oryzae* deploys various of effectors inside the rice to subvert host immunity response and facilitate host colonization [2]. Earlier studies have revealed two distinct effector secretion systems in *M. oryzae* [2, 5]. Effectors, such as LysM protein 1 and Bas4, are delivered into the apoplastic compartment where enclosed by extensions of the plant plasma membrane termed the extra-invasive hyphal membrane (EIHM), outlining the entire IH uniformly via the conventional secretion pathway [6, 7]. The conventional secretion starts at the endoplasmic reticulum (ER) and routes proteins to the Golgi apparatus via COPII-coated vesicles, and then proteins destined for the cell surface can be sorted into transport vesicles for delivery to the plasma membrane [8]. In *M. oryzae*, a novel COPII uncoating factor MoSwa2, interacts with MoSec24-2 and coordinates secretion of apoplastic effectors [9]. Additionally, the COPII cargo receptor MoErv29 plays important roles in mediating extracellular laccases and apoplastic effector secretion [10], indicating the importance of the conventional secretion pathway in secretion of apoplastic effectors.

In contrast to the conventional secretion pathway, leaderless cytoplasmic proteins can be translocated across the plasma membrane through the unconventional protein secretion (UPS) pathway [11]. To date, four distinct UPS secretion mechanisms have been identified [12, 13]. In Type I and Type II secretions, cargo proteins are translocated across the plasma membrane through pore formation or translocated through ABC transporters [11, 14, 15]. In Type III secretion, the related cargoes are sorted into autophagosomal or endosomal vesicles, which in turn fuse with the endocytic recycling compartment to empty their contents into the extracellular space in an Sso1-dependent manner [13, 16, 17]. In Type IV secretion, signal peptide-containing proteins or specific transmembrane proteins bypass the Golgi via GRASP- or HSP70-mediated transport [13, 18]. In *M. oryzae*, the cytoplasmic effectors, such as Avr-Pita, Avr-Pizt, Pwl1 and Pwl2, first accumulate in a novel membrane structure called the biotrophic interfacial complex (BIC), and are then delivered into the host cells via the exocyst complex, and MoSso1-mediated secretion [5]. Since Brefeldin A (BFA, that inhibits conventional ER to Golgi secretion) treatment had no effect on the secretion of cytoplasmic effectors into the BIC, implying that BIC-associated cells principally secrete effectors using the UPS pathway. Although the BIC has emerged as delivery site of *M. oryzae-*deployed virulence factors, how these effectors are transported to the BIC structure remains largely unclear. Thus, understanding the unconventional trafficking pathway, especially as a point for effector transport into the BIC will be essential to fully interpret the function of such intracellular processes.

Retromer, an ancient and highly conserved multimeric protein complex, is vital for regulating the retrieval and recycling of numerous cargoes away from the degradative pathway for delivery to the trans-Golgi network (TGN). In addition to its role in retrograde trafficking, an increasing body of evidence indicates that retromer is also required for endosome to cell surface recycling [19, 20]. Reflecting its central importance, retromer is essential for development and its dysfunction is linked to a number of ailments, including Alzheimer’s, and Parkinson’s disease [21, 22]. In spite of the accumulating evidence showing the importance of retromer complex in endosome–plasma membrane protein trafficking, the actual demonstration of the presence of free retromer vesicles as a transport vector at the endosome–plasma membrane interface is still elusive, especially during pathogen-host interaction.

Our previous studies in *M. oryzae* demonstrated that the retromer complex plays vital roles in retrograde trafficking, fungal development and plant infection [23, 24]. Consistent with studies in yeast, *Fusarium graminearum* and mammals, recruitment of retromer to endosomal membrane requires the Rab GTPase MoRab7 in *M. oryzae* [25–27]. Rab7 is a highly conserved protein that belongs to the Ras superfamily of small GTPase, which plays important roles in the endolysosomal system [28, 29]. The dynamic between the active GTP-bound state and inactive GDP-bound form is required for proper functioning of Rab7 [29]. Similar to retromer mutants, loss of MoRab7 impairs the vacuole fusion, autophagy and pathogenicity in *M. oryzae* [25, 30]. Since the UPS pathway is important for cargo secretion, and expanding studies have unveiled the roles of endolysosomal system in UPS pathway, we therefor hypothesize that Rab7 and retromer can function together in the UPS pathway to coordinate effector secretions in *M. oryzae*.

In this study, we focused on dissecting the role of MoRab7-retromer in effector secretion and virulence in *M. oryzae*. We showed that GFP-MoRab7 labeled vacuole closely associates with the BIC structure during plant infection. MoRab7 GTPase activity dysfunction impairs the BIC structure development and Pwl2 secretion. In addition, MoVps35 physically interacts with MoRab7 coordinate active trafficking and fusion of cargoes with the biotrophy interface. Using a chemical genetic inactivation approach (tet-off system), we were able to regulate MoVps35 activity and monitor its dynamic in effector secretion. We present evidence that the retromer complex plays important role in the secretion of apoplastic- and cytoplasmic-effectors. Furthermore, the retromer-regulated v-SNARE MoSnc1 was identified through MoVps35-GFP pull-down assays. The physical interaction with Vps35 ensures proper localization and distribution of the MoSnc1. Additionally, MoSnc1 interacts and associates with the biotrophy interface and the EIHM, and is required for proper secretion of effectors and for pathogenicity in the blast fungus. Collectively, in this study we dissect the functional role of MoRab7/Retromer/MoSnc1 cascade in effector secretion during *in planta* invasive growth of *M. oryzae*.

## Results

### MoRab7 is in close proximity with and adjacent to the BIC

In *M. oryzae-*invaded rice cells, the pathogen secretes cytoplasmic effector proteins such as Pwl2, into a specialized biotrophic interfacial complex (BIC). Interestingly, we found that a relatively prominent vacuole always flanks the BIC in the bulbous invasive hyphae (Fig. 1A, Video S1). To analyze/understand whether an unconventional secretion system is associated with such BIC-associated vacuoles during *M. oryzae-*rice interaction, we first generated an *M. oryzae* strain expressing Pwl2-mCherry (marker for BIC), together with GFP-MoRab7 (a generally acknowledged endosomal/vacuolar marker that mediates vesicular trafficking at the late endosomes) [25, 30]. During the growth of this dual-tagged strain inside rice cells, GFP-MoRab7 initially localized to distinct vesicular/punctate structures in the filamentous primary invasive hyphae (IH), while Pwl2-mCherry preferentially accumulated within the BIC at the hyphal tip (Fig. 1B and Fig. S1A). When the primary hyphae differentiated into bulbous IH, GFP-MoRab7 epifluorescence was evident in vacuolar membrane while Pwl2-mCherry showed persistent accumulation in the mature BIC (Fig. 1C and Fig. S1B-C). Strikingly, GFP-MoRab7 positive structures (vesicular or vacuolar membrane) were often adjacent to the BIC throughout the fungal colonization process, including the initial penetration stage, IH extension, and the cell-to-cell spread (Fig. 1 and Fig. S1). Additionally, three-dimensional (3D) rendered time-lapse imaging further confirmed the close proximity of GFP-MoRab7 and the BIC (Fig. 1 and Video S2-S3), thus implying a potential functional connection between MoRab7-derived membrane trafficking and fungus-host interface.

**Figure 1.**
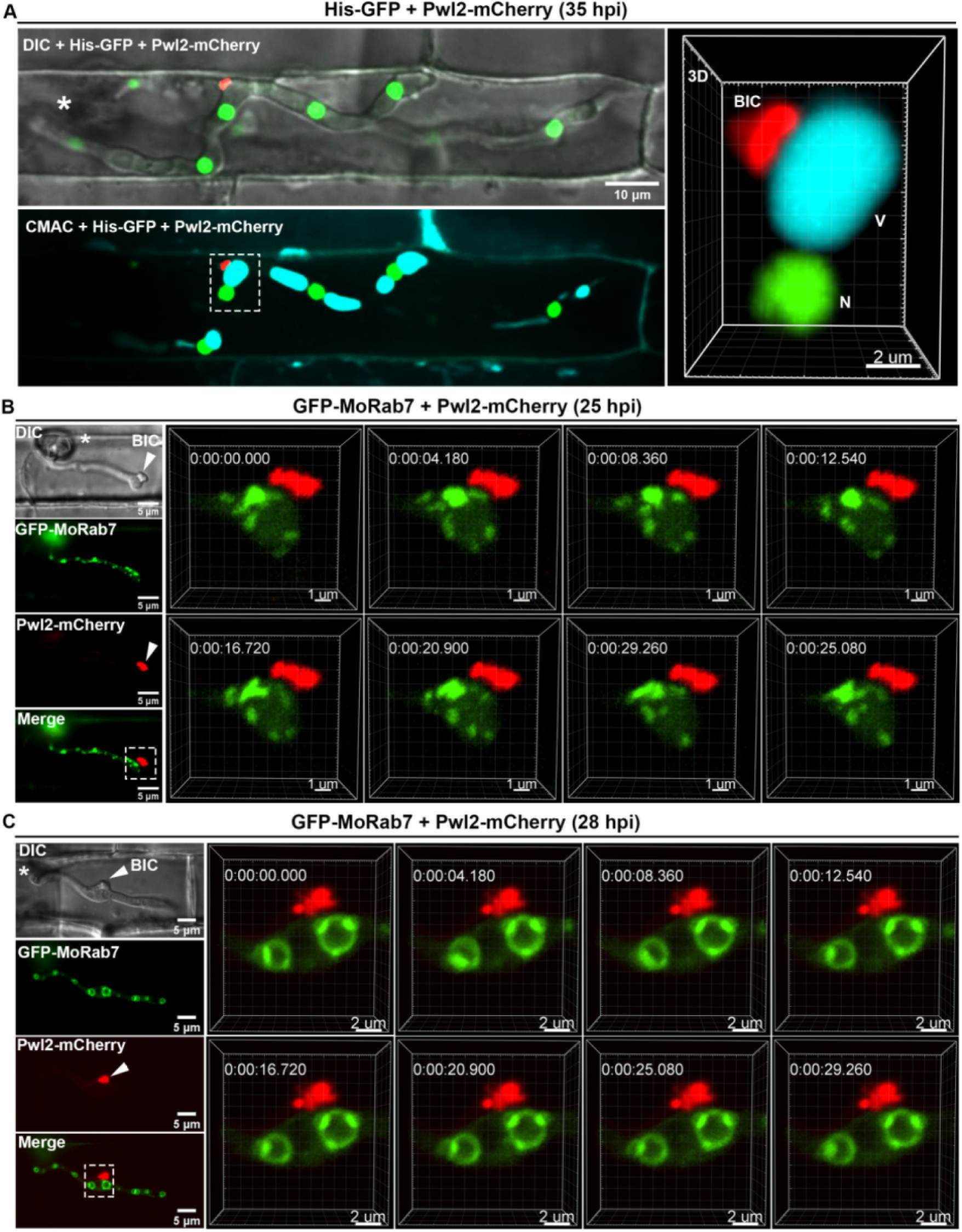
Close association of MoRab7 with the host interphase (Biotrophy Interfacial Complex; BIC) during invasion. (A) A relatively large vacuole often adjacent to the BIC during host invasion. A 3D (three-dimensional) image showing spatial positioning of the BIC, vacuole and nucleus (see also Supplementary video 1). His-GFP and Pwl2-mCherry were used to visualize the nuclei and BIC, respectively, while CMAC (7-amino-4-chloromethylcoumarin) was used to selectively stain the lumen of vacuoles. (B) At the early infection stage (25 hpi, hours post inoculation), GFP-Rab7 vesicles are abundant in the primary invasive hyphae (IH) and remain adjacent to the BIC (boxed region). Time-lapse imaging (four-dimensional) shows the dynamic trafficking/association between GFP-Rab7 and the BIC (Inset panels on the right; and Supplementary video 2). (C) At the later infection stage (28 hpi), GFP-MoRab7 mainly localizes to the vacuolar membrane in the bulbous IH adjacent to the BIC. Panels on the right depict the dynamic movement of the GFP-MoRab7 and the BIC (Supplementary video 3). DIC: differential interference contrast. Asterisks indicate appressoria. Arrowheads point to the BIC.

### MoRab7 is essential for proper BIC development but not for maintaining the apoplastic interface

Rab GTPases function as molecular switches that cycle between active (GTP-bound) and inactive (GDP-bound) states, and this cycle is linked to their membrane and cytoplasmic localization profiles [29]. In *M. oryzae,* MoRab7 null or dominant-negative (DN, a form of GDP-bound state) mutants fail to infect rice plants even when inoculated onto wounded leaves [25, 30]. To understand the role (if any) of MoRab7 in secretion of cytoplasmic effectors via BIC, we decided to conditionally inactivate MoRab7 specifically during host penetration stages of blast infection. The *PWL2* gene is specifically and strongly expressed during invasion but remains suppressed *in vitro* and during other developmental stages including vegetative growth, conidiogenesis and appressorium formation [31]. Therefore, we hypothesized that expression of an ectopic GFP-MoRab7DN construct (a form of inactive GDP-bound state) under control of the PWL2 promoter will not only ensure inherent Rab7 maintaining a normal activity before fungal penetration but also can inactivate MoRab7 due to excessive accumulation of inactive GFP-MoRab7DN within the invaded rice cells. Two independent dominant-negative alleles of Rab7 (T22N and N125I) were generated based on previous studies in yeast and mammals [25]. As expected, compared to the expression of wild-type GFP-MoRab7, the expression of GFP-MoRab7DN (T22N) or GFP-MoRab7DN (N125I) changed the localization of MoRab7 proteins to the cytoplasm in the IH (Fig. 2A-C, Fig. S2), indicating a successful inactivation of MoRab7. Under such MoRab7DN condition, Pwl2-mCherry appeared as multiple and discrete fluorescent foci in the invaded rice cells, instead of highly concentrated and specialized BIC region observed in the WT control (Fig. 2A-C, Fig. S2, video S5-S6), indicating that such developmental stage-specific inhibition of MoRab7 activity affects BIC integrity/establishment and the focal secretion of Pwl2-mCherry, leading to the production of split or distorted BICs in the MoRab7DN strains of *M. oryzae*. Next, we tested the role of a constitutively active (CA) form of MoRab7 in Pwl2 effector secretion. Our results showed that overexpression of the constitutively active GFP-MoRab7^Q67L^ variant (GTP-bound state) using PWL2 promoter does not show any defects in the vacuolar membrane localization of GFP-MoRab7 and/or the focal secretion of Pwl2 (Fig. 2D, video S7).

**Figure 2.**
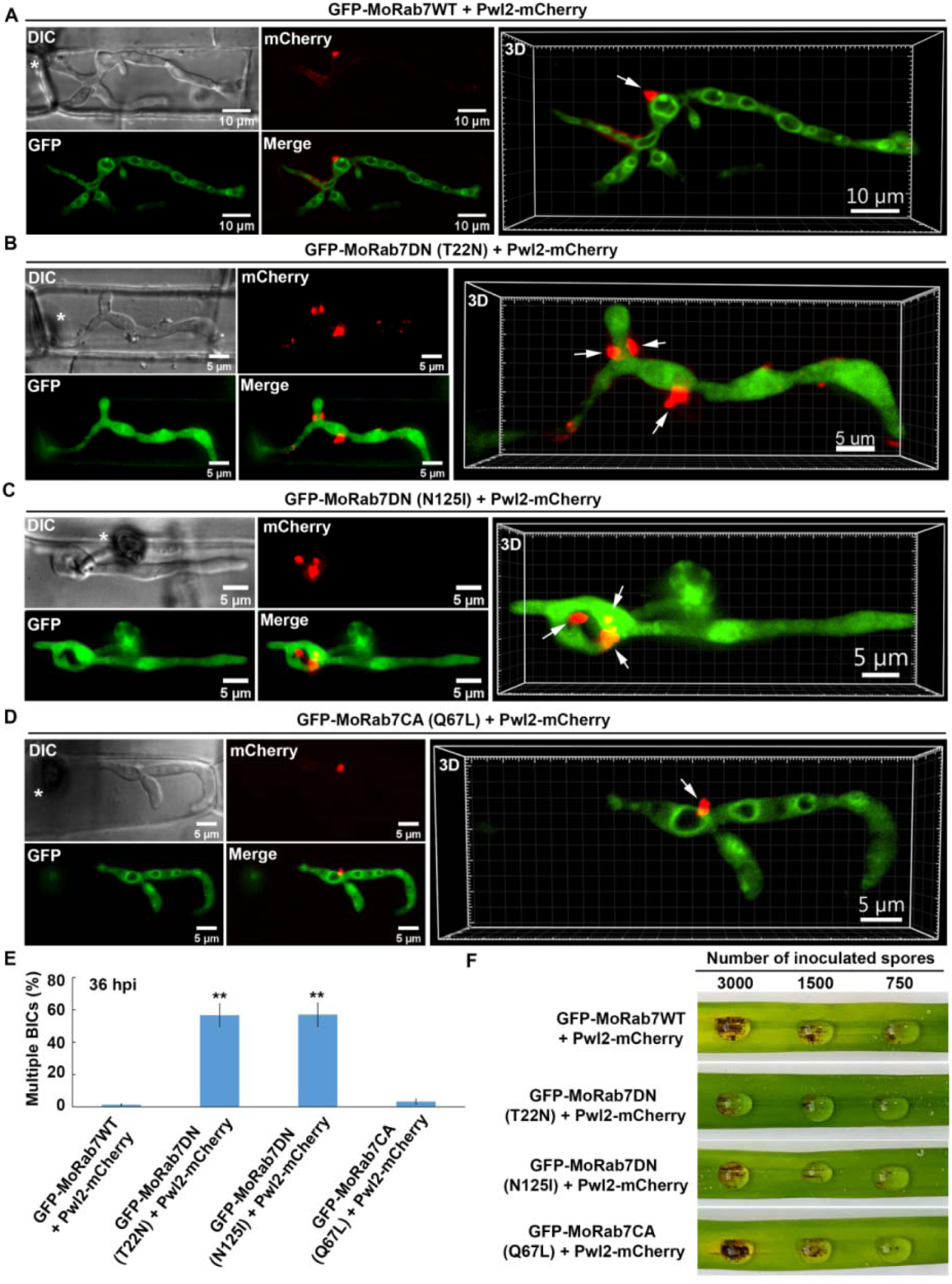
Inhibiting the GTPase activity of MoRab7 affects proper establishment of the host interface (BIC) and plant infection in the rice blast fungus. (A) Expression of GFP-MoRab7 in the wild-type (WT) under the control of *PWL2* promoter shows normal vacuolar localization, and MoRab7 is in close proximity with and adjacent to the BIC (Supplementary video 4). (B) Expression of a dominant-negative GFP-MoRab7 via T22N mutation (locked in GDP-bound state) under the control of *PWL2* promoter results in the failure of MoRab7 to target the vacuolar membrane, impairs effector secretion and produces abnormal BICs. Multiple and discrete fluorescent foci (Pwl2-mCherry, arrows) are observed in the invasive hyphae (Supplementary video 5) in the DN mutant instead of the singular and highly specialized BIC region found in the GFP-MoRab7WT. (C) Another dominant-negative mutation (N125I) of MoRab7 shows enrichment of the protein in the cytosol and causes similar aberrant secretion of Pwl2-mCherry (Supplementary video 6). (D) Constitutively active GFP-MoRab7^Q67L^ (GTP form) does not show any defects in vacuole localization and BIC development (Supplementary video 7). (E) A bar chart showing the percentage BICs formed in the IH of each of the indicated strain. (**P < 0.01; Student’s *t* test; three biological replicates; n=300 invaded cells in total). (F) Blast infection assays showing that the GTPase activity of MoRab7 is required for the fungal pathogenicity. Asterisks indicate appressoria. Arrows point to the BICs.

To understand whether the MoRab7 GTPase activity is also required for maintenance of the apoplastic interface, we used the same strategy to conditionally inactivate MoRab7 during host invasion by *M. oryzae*. The results showed that inhibition of MoRab7 GTPase activity does not affect the localization of Bas4, an apoplastic effector, during invasive growth in the rice sheath (Fig. S3). Since efficient secretion of effectors has a role in the development of rice blast disease, we next investigated the pathogenicity of GFP-MoRab7^WT^, GFP-MoRab7^DN^(T22N), GFP-MoRab7^DN^(N125I) and GFP-MoRab7^CA^(Q67L) strains on rice. Consistently, we observed that the inhibition of MoRab7 GTPase activity in invasive hyphae significantly reduced *Magnaporthe* pathogenicity on rice leaves (Fig. 2F). Taken together, these results led us to conclude that the MoRab7 GTPase activity is important for establishment and development of normal BIC during plant infection by the rice blast fungus.

### MoRab7 interacts with MoVps35 and both proteins are associate with the BIC during host invasion

Although GFP-MoRab7 positive structures are often adjacent to the BIC, and MoRab7 GTPase function is necessary for BIC development, we failed to observe a direct transportation of GFP-MoRab7 from endosomes or vacuoles to the BIC (Fig. 1, video S1 and S2). One possible explanation for this is that MoRab7 may mediate other vesicular membrane trafficking event(s) to facilitate effector secretion. Recently, we identified that the retromer CSC subcomplex is recruited by MoRab7 and is sequentially sorted by MoVps17 for effective conidiation and pathogenicity in the blast fungus. Retromer is a multi-subunit protein complex that mediates the transportation of various transmembrane proteins/receptors from endosomes to the trans-Golgi network (TGN) or plasma membrane [20]. Based on these clues, we reasoned that MoRab7 may recruit the retromer CSC to facilitate effector secretion across the BIC interface in invaded plant cells. To test this hypothesis, we first investigated the relationship between MoRab7 and the core retromer subunit MoVps35 during invasion of rice cells by *M. oryzae.* Using co-localization and Co-IP assays, we found that MoRab7 interacts physically with MoVps35 and both are positioned next to the BIC during invasive growth (Fig. 3A-B). Interestingly, live-cell imaging of BIC-associated IH dynamics clearly showed that MoVps35-GFP labeled vesicles detach from mCherry-MoRab7-labeled vacuolar membrane, and subsequently move towards/into the BIC interface over time (Fig. 3C Video S8), thus supporting the idea that the retromer CSC subcomplex is recruited by MoRab7 and sequentially sorted to the BIC-associated zone in *M. oryzae*.

**Figure 3.**
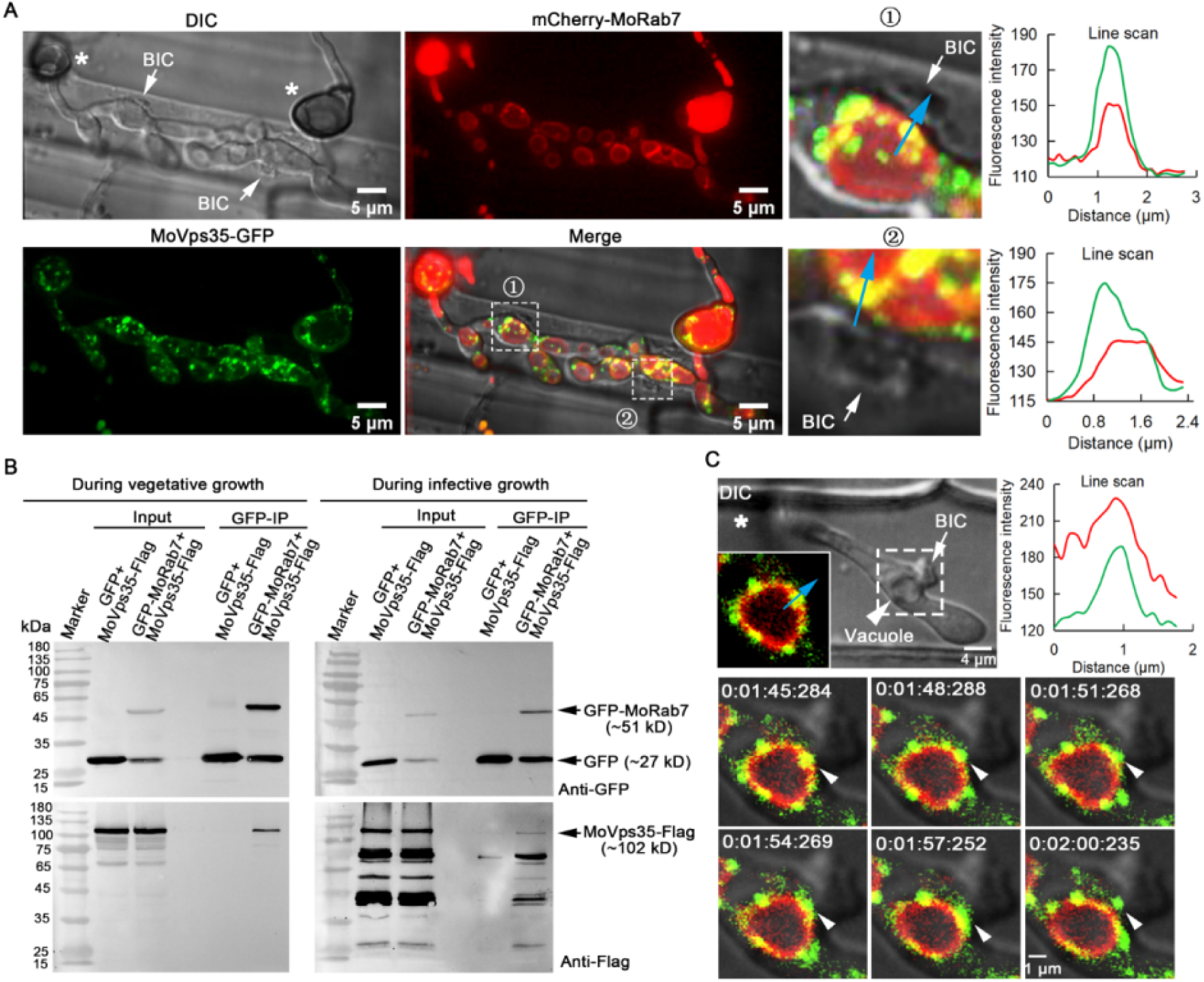
MoRab7 interacts with retromer MoVps35 and both are positioned next to the BIC during host invasion. (A) A projection image of infectious hyphae of the strain co-expressing mCherry-MoRab7 and MoVps35-GFP. The mCherry-MoRab7 localizes to the vacuolar membrane in the bulbous IH, while MoVps35-GFP is mainly localized to punctate structures that surround or attach to the vacuolar membrane. In the detailed view of the regions ① and ②, the mCherry-MoRab7 and MoVps35-GFP partially co-localized (yellow) adjacent to the BIC. Blue arrows in the insets show the path for fluorescence intensity distribution by line-scan analysis. (B) GFP-trap-based pull-down experiment indicating the physical interaction between MoRab7 and MoVps35 during the fungal vegetative and *in planta* growths. (C) Time-series montage showing the co-localization of mCherry-MoRab7 and MoVps35-GFP near the BIC region. The highlighted region shows the dynamic trafficking of MoVps35-GFP (arrowhead) from the vacuolar membrane to the BIC interface over time (Supplementary video 8). The vacuoles appear hollow in the DIC image. Asterisks indicate appressoria. White arrows point to the BICs.

### Retromer complex mediates membrane trafficking to the BIC and the EIHM during host invasion

To analyze the details of retromer-mediated trafficking across the fungus-plant interface, we co-expressed MoVps35-GFP (the core subunit of the retromer) and MoVps17-GFP (the key sorting nexin of the retromer), together with Pwl2-mCherry, respectively. In the resulting transformants, both MoVps35-GFP and MoVps17-GFP localized to distinct punctae, some of which were clearly adjacent to the Pwl2-mCherry-decorated BIC during invasive growth of *M. oryzae* (Fig. 4A-E, Fig. S4A). Using surface rendering, we reconstructed the epifluorescent interface between Pwl2-mCherry and MoVps35-GFP or MoVps17-GFP (Fig. 4A, 4B, 4D). This analysis clearly showed the intimate positioning of the retromer vesicular sorting machinery (MoVps35-GFP or MoVps17-GFP) and the BIC (Pwl2-mCherry) at the interactive interfaces (Fig. 4A, 4B, 4D, Video S9 and S10). In addition, time-lapse imaging showed that MoVps35-GFP- and MoVps17-GFP-labelled vesicles move to the BIC-associated zone and later fade away (Fig. 4C and 4E, video S11 and S12), implying a likely fusion with the target membrane. We also analyzed the dynamics between MoVps35-GFP and Pwl2-mCherry in invasive hyphae in plasmolyzed rice cells, since sucrose-induced plasmolysis concentrates BIC-associated membranes [32]. The results showed that MoVps35-GFP-labeled vesicles were still localized to the IH region adjacent to the BIC, thus displaying a sorting and fusion nature with the target membrane underneath the BIC (Fig. S5, video S13). However, we could not detect any direct interaction between MoVps35 and Pwl2 by bimolecular fluorescence complementation (BiFC) assay (Fig. S7). Next, we investigated the spatiotemporal dynamics of MoVps35-GFP with the apoplastic effector Bas4-mCherry which accumulates in the EIHM outlining the invasive hyphae during *M. oryzae* infection. We observed that several MoVps35-GFP punctae adjacent to the EIHM throughout the fungal colonization process (Fig. 4F, Fig. S7 and video S14). Consistently, time-lapse imaging further confirmed that MoVps35-GFP displays a dynamic fusion process on the plasma membrane of the invasive hyphae (Fig. 4G, video S15). We infer that the retromer complex is involved in vesicular trafficking from endosomes to the plasma membrane, and the BIC-specific membrane during *M. oryzae* pathogenesis.

**Figure 4.**
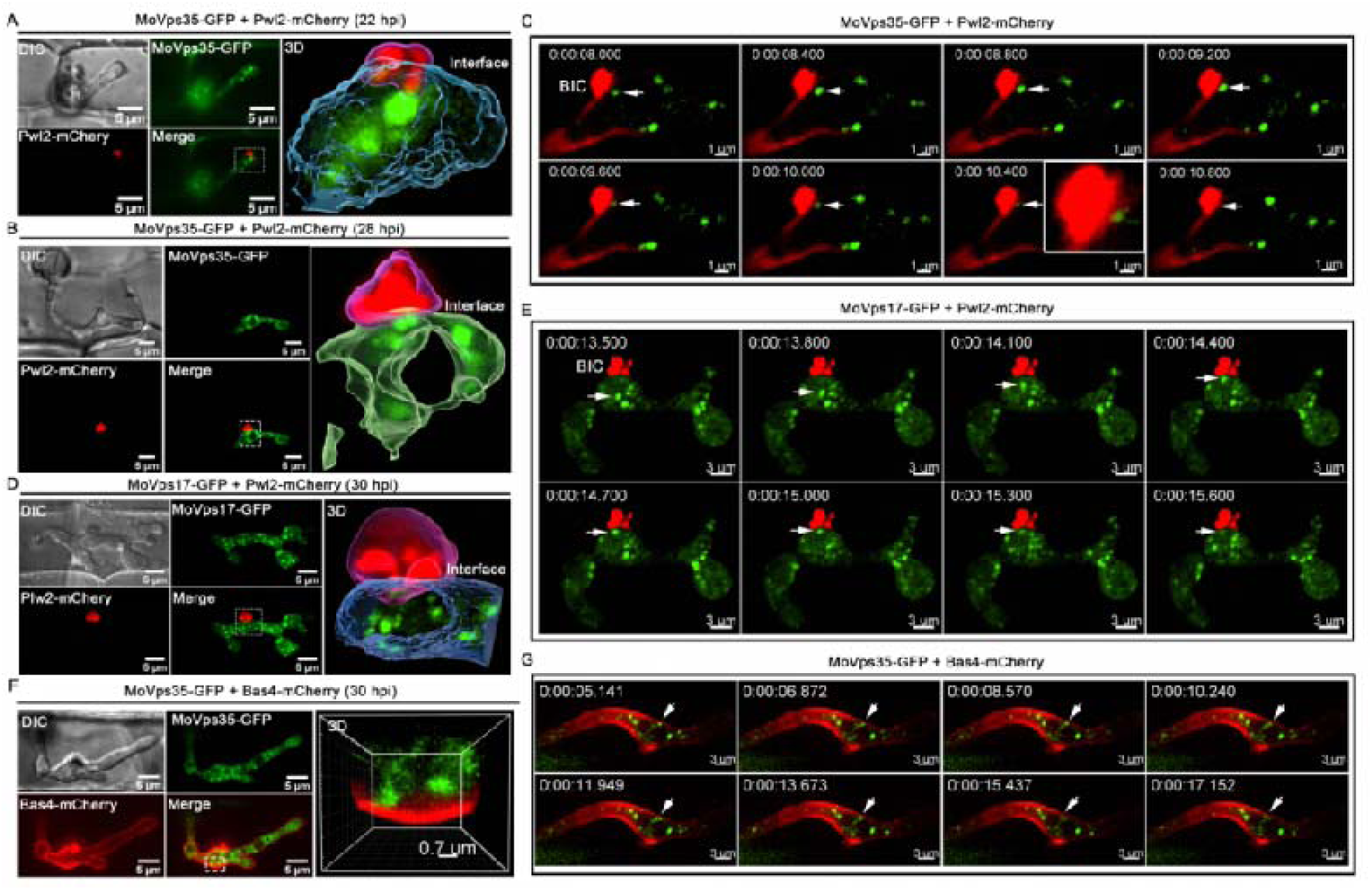
Association of the endosomal sorting machinery, retromer complex, with BIC and EIHM during host invasion. (A) Confocal micrographs of filamentous primary invasive hyphae expressing MoVps35-GFP (an indispensable retromer subunit) and Pwl2-mCherry (BIC-specific marker) at 22 hpi. At the hyphal tip, the retromer is distinctly adjacent to the BIC (see boxed region). The 3D image (right panel) positioning of the vesicular cargo sorting machinery (MoVps35-GFP) and the BIC (Pwl2-mCherry) at the host interface (Supplementary video 9). Image processing was performed using Imaris 9.5 with the surface rendering tool. (B) At 28 hpi, the filamentous primary IH develop into bulbous hyphae wherein MoVps35 localizes to the vacuolar membrane adjacent to the BIC (Supplementary video 10). (C) Time-lapse confocal imaging of the MoVps35-GFP and Pwl2-mCherry in the bulbous infection hyphae. Notably, a MoVps35-GFP-labelled vesicle (diameter ≤ 0.5 μm; arrow) in the IH moved towards and/or transiently fused with the BIC-associated interface (Supplementary video 11). The instant contact point is depicted in the detailed view (see time 0:00:10:400). Images are shown at 400 millisecond intervals. (D) Visualization of infectious hyphae expressing MoVps17-GFP (the retromer-associated sorting nexin) and Pwl2-mCherry at the bulbous IH (30 hpi). MoVps17-GFP also localizes to distinct vesicles that are clearly adjacent to the BIC (see dotted box). The 3D rendered image (right panel) shows the retromer sorting machinery (MoVps17-GFP) proximal to the BIC (Pwl2-mCherry) at the interface. (E) Time-lapse confocal microscopy of the bulbous invasive hyphae of the MoVps17-GFP Pwl2-mCherry strain. Consistent with MoVps35-GFP, the MoVps17-GFP-labelled vesicle (arrow) moves dynamically to the BIC-associated plasma membrane of the invasive hyphae (Supplementary video 12). Images are shown at 300-ms intervals. (F) The retromer also associates with the sites of secretion of the apoplastic effectors. Micrographs depicting the localization of MoVps35-GFP and Bas4-mCherry (EIHM/apoplast-specific marker) within the bulbous IH (30 hpi). MoVps35-GFP localizes to some distinct punctate structures, some of which are clearly adjacent to the EIHM (see boxed region). The 3D rendition (right panel) shows an intimate positioning of the MoVps35-GFP and the Bas4-mCherry marked EIHM (Supplementary video 14). (G) Time-lapse confocal imaging of the bulbous infection hyphae expressing MoVps35-GFP and Bas4-mCherry. A MoVps35-GFP-labelled vesicle (arrow) in the IH was clearly observed moving to the EIHM and gradually fades away (Supplementary video 15).

### The retromer complex is essential for the secretion of apoplastic and cytoplasmic effectors in *M. oryzae*

*ΔMovps35* mutant does not penetrate the rice cuticle efficiently because of the lack of retromer function during autophagy-dependent plant infection [23]. However, few penetration events by the *ΔMovps29* mutant were evident despite a delay and reduction in the ability of the mutant to spread between rice cells (Fig. 5A). We therefore co-expressed Bas4-GFP and Pwl2-mCherry in the *ΔMovps29* mutant, together with the wild type, and investigated their secretion during *M. oryzae*-rice interaction. We found that Bas4-GFP mostly accumulated inside the vacuoles and Pwl2-mCherry appeared as multiple and discrete epifluorescent foci in the *ΔMovps29* mutant invasive hyphae (Fig. 5), suggesting that the disruption of the retromer complex impairs the secretion of both the apoplastic and the cytoplasmic effectors in the blast fungus.

**Figure 5.**
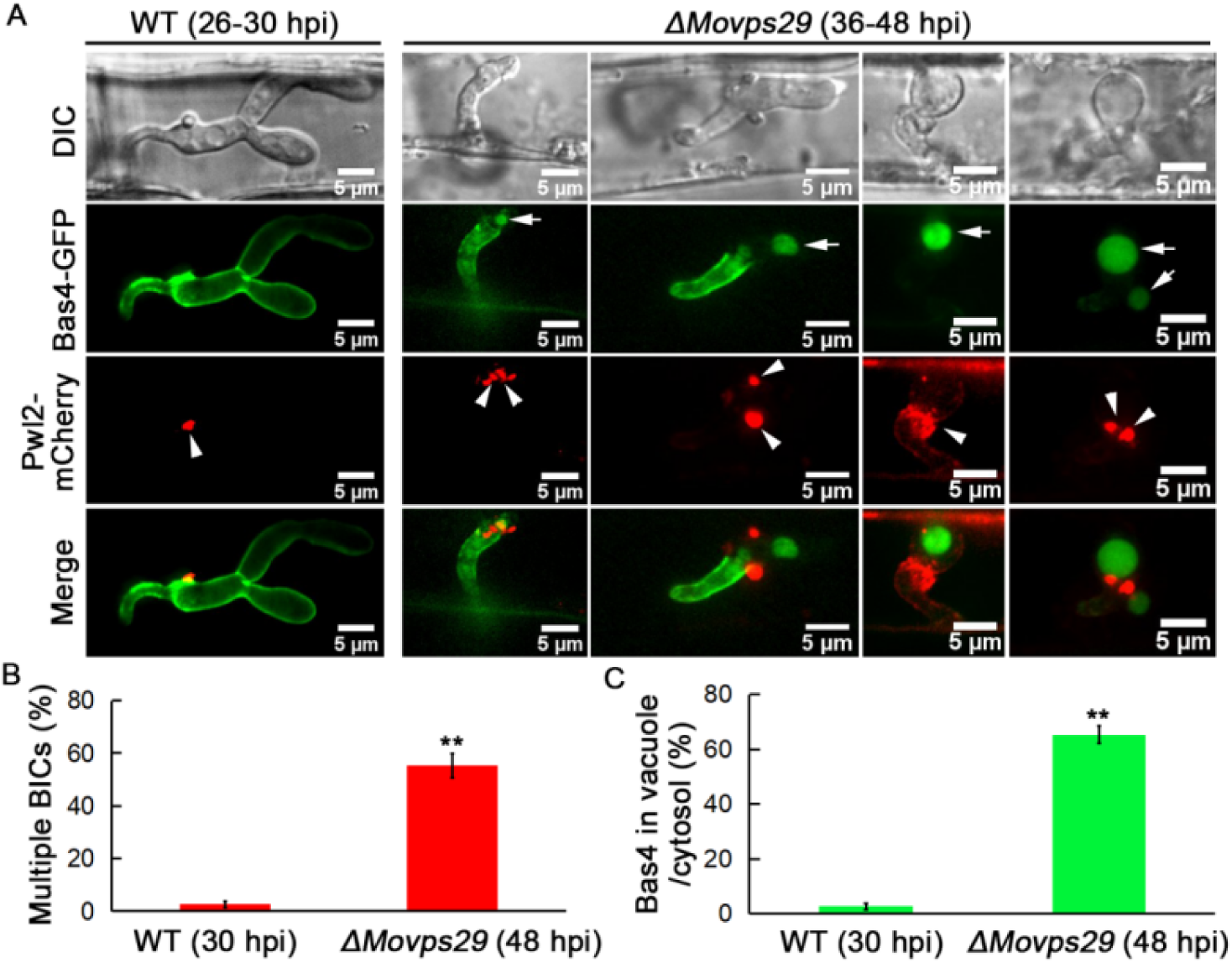
Disruption of the retromer complex impairs secretion of both apoplastic and cytoplasmic effectors. (A) Bas4-GFP (apoplastic) and Pwl2-mCherry (cytoplasmic) were improperly secreted and mislocalized in the *ΔMovps29* mutant. In the WT, Bas4-GFP shows apoplastic localization, outlining the invasive hyphae, while it often accumulates inside the vacuole (arrows) in the *ΔMovps29* mutant. Pwl2-mCherry appeared as multiple and discrete fluorescent foci (arrowheads) in the *ΔMovps29* invasive hyphae, instead of focussed and specialized BIC observed in the WT. (B) A bar chart showing the percentage of IH containing multiple BICs in the WT and *ΔMovps29* mutant. (C) A bar chart showing the percentage of IH containing vacuolar Bas4 in the indicated strains. (**P < 0.01; Student’s *t* test; three biological replicates; n=150 invaded cells in total).

### Inactivation of MoVps35 impairs effector secretion in *M. oryzae*

To confirm that the retromer complex is important for effector secretion during *M. oryzae*-rice interaction, we decided to conditionally inactivate the MoVps35 using a chemical genetic approach. We established a tetracycline-regulated *MoVPS35* expression system by inserting a Tet-Off cassette downstream of the *MoVPS35* promoter (Fig. S8A-D). Tetracycline-regulated gene expression in fungi such as *S. cerevisiae*, *C. albicans*, *A. fumigatus* and *Ustilago maydis* has been widely applied [33–36], but not yet to elucidate the performance of the system during plant infection. Expression of the tetracycline-regulated *GFP-MoVPS35* gene under the control of its native promoter displayed punctate localization of the protein on endosomal membrane, consistent with the localization of MoVps35-GFP in the complemented strain (Fig. S9E). Addition of tetracycline or doxycycline selectively inhibited the expression of GFP-MoVps35 during vegetative/mycelial growth (Fig. S9F). Furthermore, treatment of Tet-Off-GFP-MoVps35/Pwl2-mCherry strain with doxycycline delayed the mobilization of cytoplasmic content into the appressoria, and compromised lesion development on inoculated barley leaves, similar to the defects observed in *MoVPS35* deletion mutant (Fig. S9A-C, [23]). We also observed that the addition of doxycycline to conidia of the Tet-Off-GFP-MoVps35/Pwl2-mCherry strain resulted in a significant decrease in appressorium-mediated host penetration at 28 hpi and delayed invasive growth and development at 40 hpi (Fig. S9D-F). An abnormal distribution of Pwl2-mCherry was also evident in these slow-growing invasive hyphae (Fig. S9D and S9G). To rule out the possibility that the defect in IH development may be the cause of the abnormal secretion of Pwl2 effector, we allowed the Tet-Off-GFP-MoVps35/Pwl2-mCherry strain to invade the first rice epidermal cell before adding the doxycycline at 18 hpi. This treatment does not affect the frequency of appressorium-mediated host penetration but turns off the expression of *MoVPS35* specifically in the invasive hyphae (Fig. 6A), creating a suitable condition to study effector secretion in the infected rice cells. These results demonstrate that the conditional inactivation or knock-down of MoVps35 causes Pwl2-mCherry to appear as multiple and discrete fluorescent foci in the IH (Fig. 6A), suggesting an impaired and abnormal BIC and impaired Pwl2 effector secretion in *M. oryzae.* Next, we used the same strategy, Tet-Off, to conditionally inactivate the MoVps35 to investigate the secretion of Bas4 during plant invasion by the pathogen (Fig. S10). The results showed that the selective turn off of *MoVPS35* expression during invasive growth leads to mislocalization of the Bas4-mCherry effector into the vacuoles or cytoplasm of the invasive hyphae (Fig. 6B and Fig.S10F). We conclude that the retromer complex functions in mediating effective and proper secretion of cytoplasmic and apoplastic effectors during *M. oryzae*-rice interaction.

**Figure 6.**
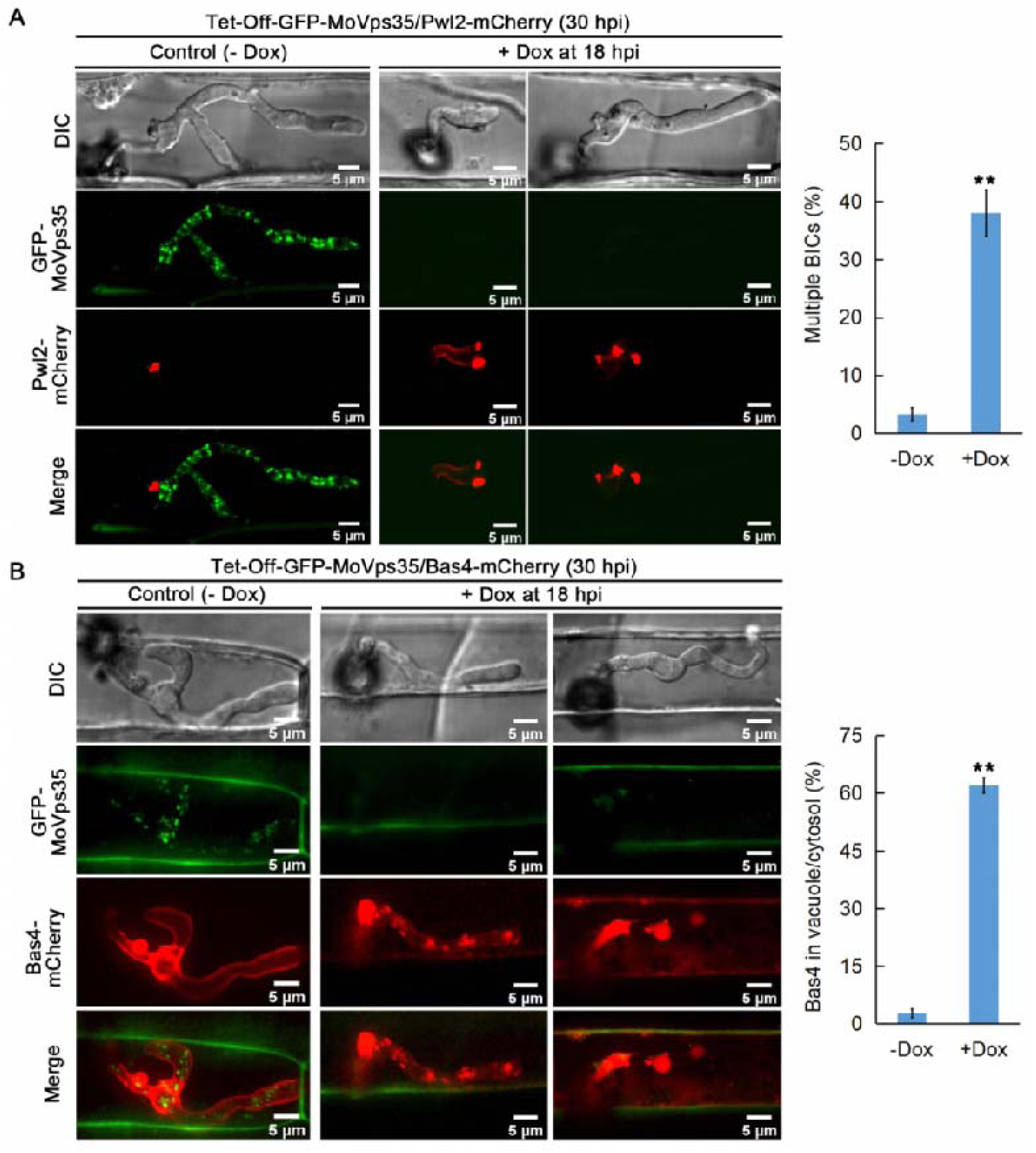
Conditional inactivation of retromer/*MoVPS35* function impairs interface establishment and effector secretion in *M. oryzae.* (A) Addition of Dox to the Tet-Off-GFP-MoVps35/Pwl2-mCherry strain blocked MoVps35 expression and results in a significantly increased number of BICs in the invasive hyphae. The bar chart shows the percentage of IH containing multiple BICs at 30 hpi after treatment with 50 µg/ml Dox at 18 hpi (**P < 0.01; Student’s *t* test; three biological replicates; 150 infected cells observed). (B) Addition of Dox to the Tet-Off-GFP-MoVps35/Bas4-mCherry strain blocked MoVps35 expression and concurrently caused Bas4-mCherry effector miss-localization into the vacuole or cytosol of the IH. The bar chart shows the percentage of IH containing vacuolar/cytosol Bas4 at 30 hpi after treatment with 50 µg/ml Dox at 18 hpi (**P < 0.01; Student’s *t* test; three biological replicates; 150 infected cells observed).

### The MoVps35-MoSnc1 interaction is required for v-SNARE based fusion of secretory vesicles with the plasma membrane

To investigate how retromer complex regulates secretion/targeting of effectors to the plasma membrane, we immunoprecipitated MoVps35-GFP and performed liquid chromatography-tandem mass spectrometry (LC-MS/MS) to identify the binding partners. Interestingly, we found several SNARE proteins to be highly enriched in the MoVps35-GFP immunoaffinity purification assays (Table 1, Table S1). *M. oryzae* genome encodes 22 SNARE proteins, whereas 12 SNARE thereof are predicted to be associated with the retromer complex, including previously identified Syn8, which is involved in the secretion of cytoplasmic effectors [37]. We noted that MoSnc1 (MGG_12614) was the most abundant amongst these 12 SNAREs (MoVps35 interactors) and was repeatedly found. In *Fusarium graminearum*, FgSnc1 is involved in regulating the fusion of secretory vesicles with the plasma membrane [38]. In *M. oryzae*, MoSnc1 was previously shown to accumulate at the hyphal growth zones and also in subapical BIC-associated cells [5], but its function has remained largely unexplored. In addition, the functional association between the retromer complex and MoSnc1 has not been identified previously. To confirm MoVps35 association with MoSnc1 *in vivo*, we generated a strain that co-expresses GFP-MoSnc1 and MoVps35-Flag and subjected it to requisite immunoprecipitation and subsequent immunoblot analysis during vegetative and invasive growth in *M. oryzae*. A 40.5 kD band corresponding to GFP-MoSnc1 and a 102 kD band corresponding to MoVps35-Flag were identified (Fig. 7A), suggesting a positive *in vivo* interaction between the MoVps35 and MoSnc1. Live-cell imaging of *M. oryzae* strain that co-expresses MoVps35-GFP and mCherry-MoSnc1 showed partial co-localization pattern as punctate structures in vegetative hyphae, conidia and invasive hyphae (Fig. 7B). In addition to localization as punctate structures, mCherry-MoSnc1 was also present at the plasma membrane and septum in basal hyphae, while actively accumulated at the apex in growing hyphal tips (Fig. 7C, Fig. S11 and video S16), suggestive of the conserved roles of MoSnc1 in regulating the fusion of secretory vesicles with the plasma membrane. Furthermore, we found that deletion of *MoVPS35* perturbed the transport of MoSnc1 to the plasma membrane due to mis-sorting of MoSnc1 to the degradation pathway (Fig. 7C). These data established that the retromer complex is able to recognize MoSnc1 at the endosomal membranes and facilitates its traffic in regulating the fusion of secretory vesicles with the plasma membrane.

**Table 1.**
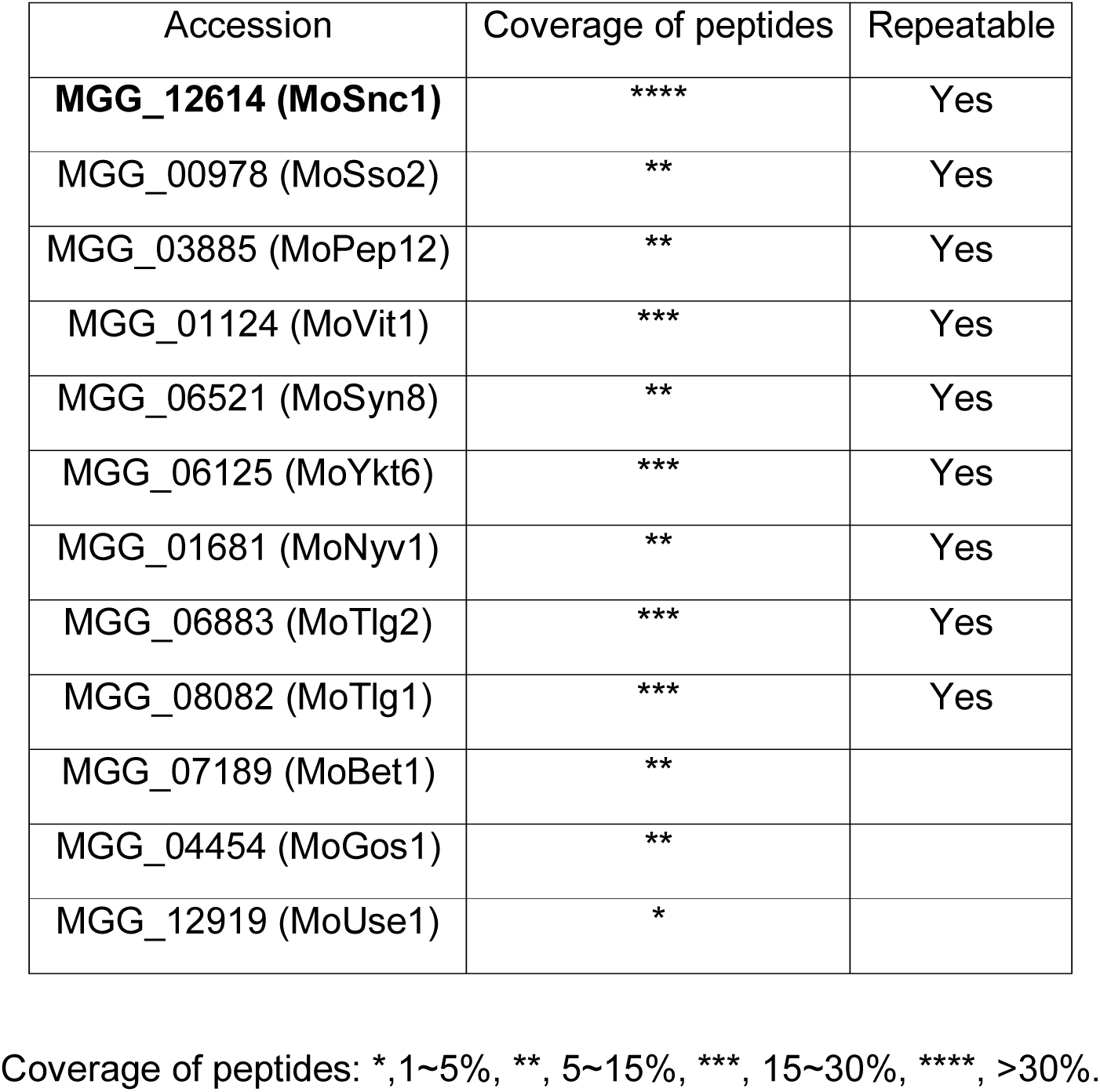
Putative SNARE interactors for MoVps35–GFP

**Figure 7.**
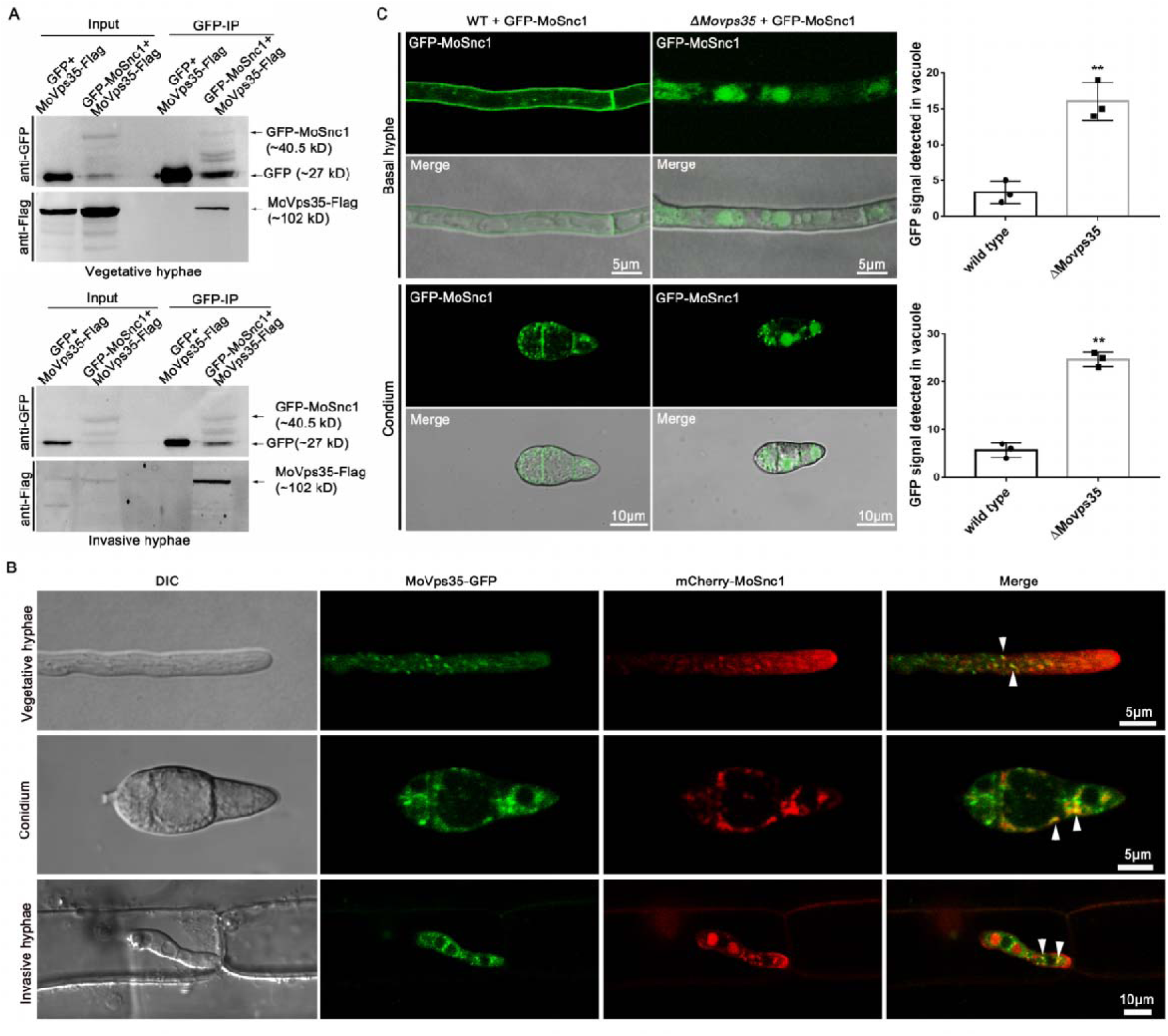
MoVps35 interacts with the v-SNARE MoSnc1 and the association is required for MoSnc1 recycling to the plasma membrane. (A) GFP-trap-based pull-down experiment indicating interaction between MoVps35 and MoSnc1 during *M. oryzae* vegetative and *in planta* growths. The strains co-expressing the indicated proteins were immunoprecipitated with GFP-trap beads. The IP signal (GFP-MoSnc1) and the Co-IP signal (MoVps35-Flag) were detected by immunoblotting with GFP and Flag antibodies, respectively. (B) Representative confocal micrographs showing partial co-localization (in yellow; Arrowheads) between MoVps35-GFP and mCherry-MoSnc1 in vegetative hyphae, conidia and invasive hyphae. (C) Deletion of *MoVPS35* perturbs MoSnc1 transport to the plasma membrane by mis-sorting MoSnc1 towards the degradation compartments. In the WT hyphae or conidia, GFP-MoSnc1 was detected on the plasma membrane. In contrast, GFP-MoSnc1 was missorted into the vacuoles in the *ΔMovps35* mutant. The bar charts show the frequency of GFP-MoSnc1 in the vacuoles of hyphae and conidia in the wild type or the *ΔMovps35* mutant. (**P < 0.01; Student’s *t* test; three biological replicates; n=30).

### MoSnc1 associates with the BIC and EIHM during *M. oryzae*-rice interaction

We further investigated MoSnc1 dynamics with the Pwl2 and Bas4 during *M. oryzae*-rice interaction, respectively. At the initial penetration stage (before 22 hpi), the GFP-MoSnc1 signal was observed at the tips of filamentous IH while Pwl2-mCherry signal was not visible at this time point, suggesting that MoSnc1 expression and localization precedes that of Pwl2 (Fig. S12A). Subsequently, the GFP-MoSnc1 remained concentrated/confined at the invasive hyphal tip, concomitant with the appearance of the Pwl2-mCherry (Fig. 8A, Fig. S12A), and the two signals remained in very close proximity to each other (Fig. 8A). Consistent with the observations in the IH co-expressing MoVps35-GFP and Pwl2-mCherry, the GFP-MoSnc1-labelled vesicles in the IH too moved towards the BIC and later faded away (Fig. 8B, video S17), implying a likely fusion with the plasma membrane. Remarkably, GFP-MoSnc1-labelled small vesicles trafficked out from the IH and were delivered into the BIC (Fig. 8C, video 18), suggesting that MoSnc1 could be mediating an inter-membrane trafficking from endosomes to the BIC. In addition, we confirmed that some of the GFP-MoSnc1-labelled punctae were clearly adjacent to the EIHM throughout the entire *M. oryzae* colonization process (Fig. 8D, Fig. S13, videos S19 and S20). These data led us to conclude that MoSnc1 associates with the BIC and EIHM during *M. oryzae*-rice interaction.

**Figure 8.**
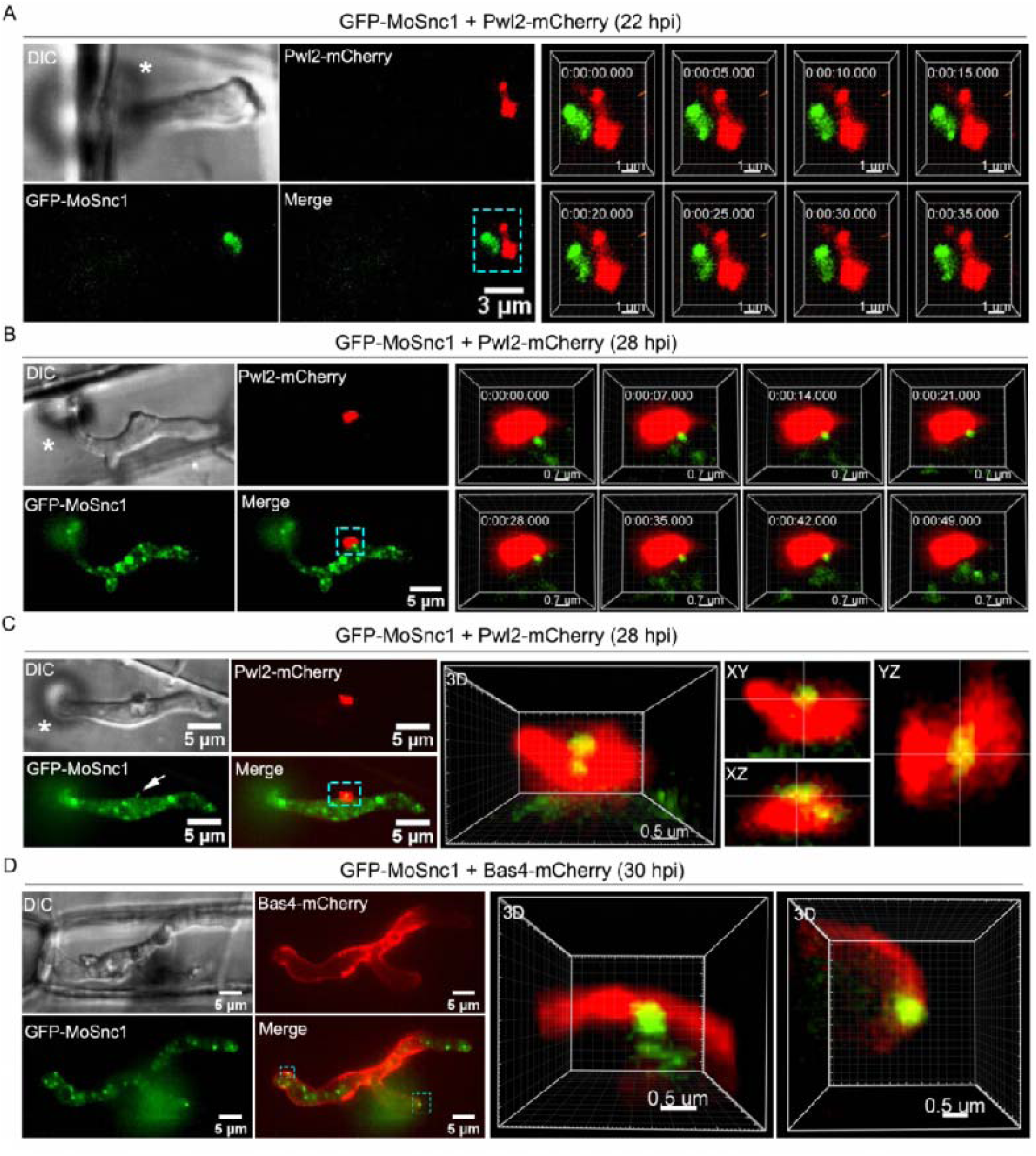
The v-SNARE MoSnc1 associates with the BIC and EIHM during host invasion. (A) Confocal micrographs of the primary invasive hyphae (22 hpi) expressing GFP-MoSnc1 and Pwl2-mCherry. At the hyphal tip, the concentrated GFP-MoSnc1 fluorescence is clearly adjacent to the tip-localized Pwl2-mCherry. The right panel shows a 3D reconstruction of the optical sections showing close association of the GFP-Snc1 and the tip BIC (Supplementary video 17). (B) Confocal image showing an intimate association between GFP-MoSnc1 and the BIC in bulbous hyphae *in planta* (28 hpi). Right panel, a GFP-MoSnc1-labelled vesicle in the IH is trafficked to the BIC and later fades away (Supplementary video 18). (C) A confocal image showing the delivery of GFP-MoSnc1 vesicles to the BIC. Notably, two GFP-MoSnc1-labelled small vesicles with diameters ≤ 0.5 μm (arrow) are trafficked out of the invasive hypha and delivered into the BIC (see the zoom in view or the 3D Supplementary video 19). The orthogonal views XY, XZ and YZ clearly illustrate co-localization of GFP-MoSnc1 and Pwl2-mCherry. Co-localization is visible as yellow fluorescence. (D) Visualization of bulbous IH (30 hpi) expressing GFP-MoSnc1 and Bas4-mCherry (Apoplast/EIHM-specific marker). Some of GFP-MoSnc1-labelled puncta are clearly adjacent to the EIHM (see boxed region). The 3D rendering (right panel) shows an intimate positioning of the GFP-MoSnc1 and the EIHM (Bas4-mCherry) (Supplementary video 20-21). Asterisks indicate appressoria.

### The v-SNARE protein MoSnc1 is required for effector secretion and fungal pathogenicity in rice blast

To understand the role of MoSnc1 in effector secretion and fungal virulence, we generated *MoSNC1-*deletion mutants (Fig. S14). Phenotypic analyses on vegetative growth, conidiation and appressorium formation of the *ΔMosnc1* mutant did not show any large differences compared to the WT (Fig S15). Subsequently, the cytoplasmic Pwl2-mCherry and the apoplastic effector Bas4-GFP were co-expressed in the *ΔMosnc1* mutant. Like the retromer complex component mutants, the Bas4-GFP and the Pwl2-mCherry proteins showed aberrant secretion and mislocalized at all stages of host penetration (Fig. 9A). The increased frequency of abnormal distribution of Pwl2 in split BIC, and the mislocalization of Bas4 to the vacuole/cytosol were consistently observed (Fig. 9A). Additionally, *ΔMosnc1* mutant showed significant pathogenicity defects in rice and barley seedlings, whether by spray or by conidia suspension droplet inoculation (Fig. 9B-E). Reintroduction of *MoSNC1* into the *ΔMosnc1* mutant restored the full virulence of the mutant strain (Fig. 9B-E). Therefore, we conclude that MoSnc1 is critical for efficient secretion of vesicles/effectors via the cytoplasmic and apoplastic interface(s) (BIC and EIHM, respectively) and is essential for fungal pathogenicity during the devastating blast disease in rice.

**Figure 9.**
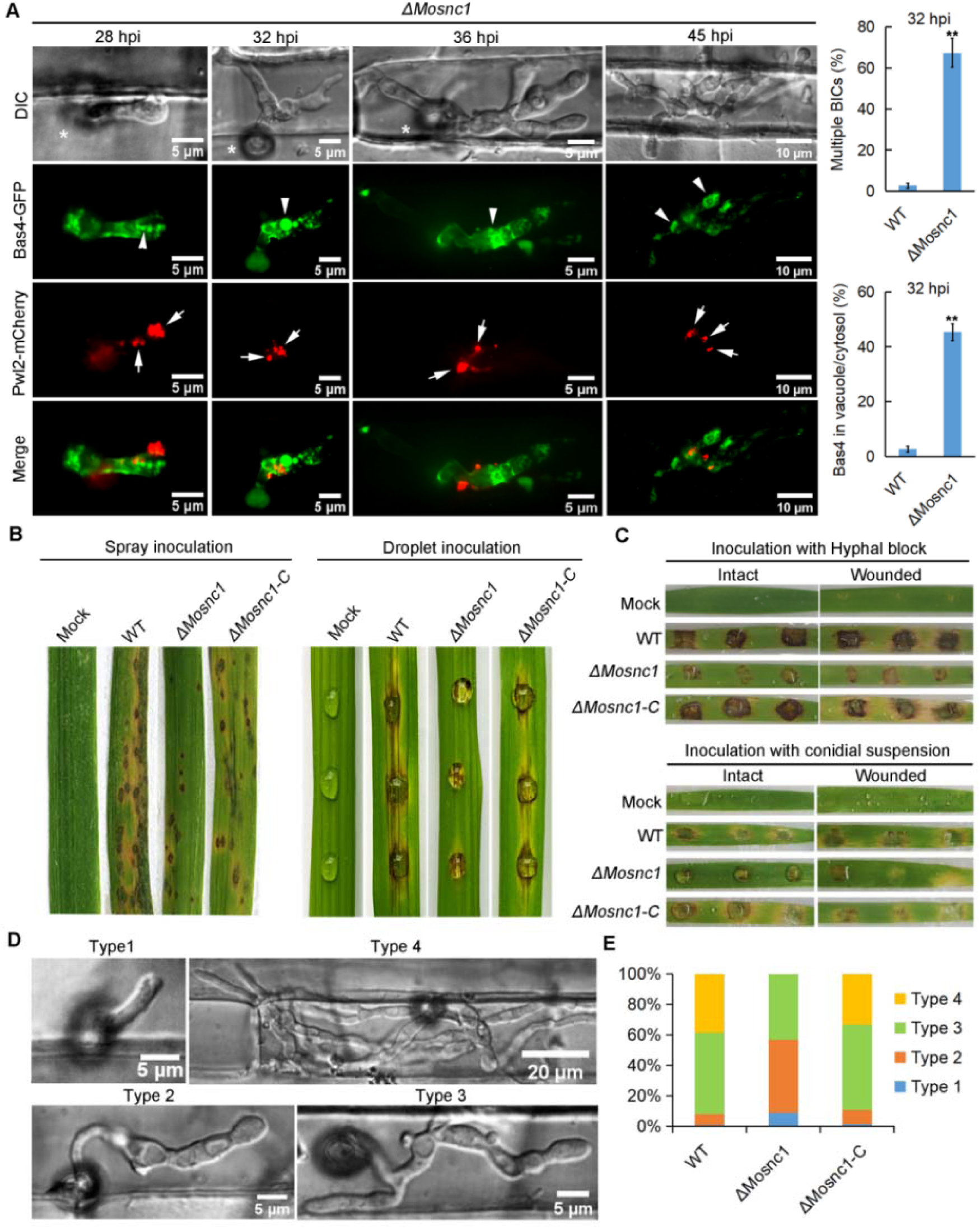
The v-SNARE protein MoSnc1 mediates proper effector secretion and fungal pathogenicity in the rice blast fungus. (A) The apoplastic effector Bas4-GFP and the cytoplasmic effector Pwl2-mCherry expressed in the *ΔMosnc1* mutant are inappropriately secreted and mislocalized at all penetration stages. In the WT, Bas4-GFP shows apoplastic localization, outlining the IH, while it often accumulates inside the vacuole of the IH (arrowhead) where it loses its outlining appearance in the *ΔMosnc1* mutant. Deletion of *MoSNC1* also causes Pwl2-mCherry fluorescence to appear as multiple and discrete fluorescent foci (arrows) in the IH of the mutant instead of one concentrated fluorescence at the specialized BIC region evident in the wild type. The bar charts show the percentages of IH containing multiple BICs and IH containing Bas4-mCherry in vacuole/cytosol in the WT and Δ*Mosnc1* mutant. (**P < 0.01; Student’s *t* test; three biological replicates; 150 infected cells observed). (B) Targeted deletion of the *MoSNC1* gene resulted in a significant reduction of *M. oryzae* pathogenicity to rice seedlings. The rice leaves (*Oryza sativa* cv. CO39) were sprayed with conidia from the wild-type strain, *ΔMosnc1* mutant and the complemented *ΔMosnc1* strain. (C) Similarly, pathogenicity defects were observed after inoculation of *ΔMosnc1* mutants on barley leaves. The barley leaves (intact and wounded) were inoculated with mycelial plugs (top panel) and conidia suspension (bottom panel). (D and E) Penetration assays in rice leaf sheath. IH growth in rice cells was observed at 38 hpi, and 4 types of IH were quantified and statistically analyzed (type 1: filamentous primary IH; type 2: a single bulbous IH; type 3: branched bulbous IH but limited to one cell; type 4: IH spread to adjacent cells). Error bars represent standard deviations. A total of 50 invaded cells were analyzed, and the experiment was repeated thrice.

## Discussion

Phytopathogenic fungi secrete various types of effectors into the host tissue to suppress plant immunity, and promote colonization and disease [2, 3]. For the translocation of the effectors, *M. oryzae* possesses two distinct secretion pathways. The apoplastic effectors are secreted via the conventional secretion system, while the cytoplasmic effectors first accumulate at the biotrophic interfacial complex and then translocated into the host cells in an exocyst and MoSso1-dependent manner [5]. Since efficient effector secretion is critical for fungal colonization, understanding such trafficking mechanism(s) would provide new insights in the development of sustainable and broad-spectrum strategies for disease control. In this study, we provided evidence that *M. oryzae* possesses a novel effector secretion pathway mediated by the MoRab7 GTPase, the retromer complex, and the associated v-SNARE MoSnc1 (Fig. 10). During invasive growth, MoRab7 is activated and then recruits the retromer complex to the vacuolar membrane to enable the proximity of retromer to the BIC structure. After attachment to the vacuolar membrane, the retromer forms a complex with and delivers the v-SNARE MoSnc1 to the plasma membrane. After reaching the plasma membrane, MoSnc1 functions as the membrane fusion catalyst that ensures the proper vesicle docking and release process required for establishment of the biotrophy associated interface/conduit for effector secretion.

**Figure 10.**
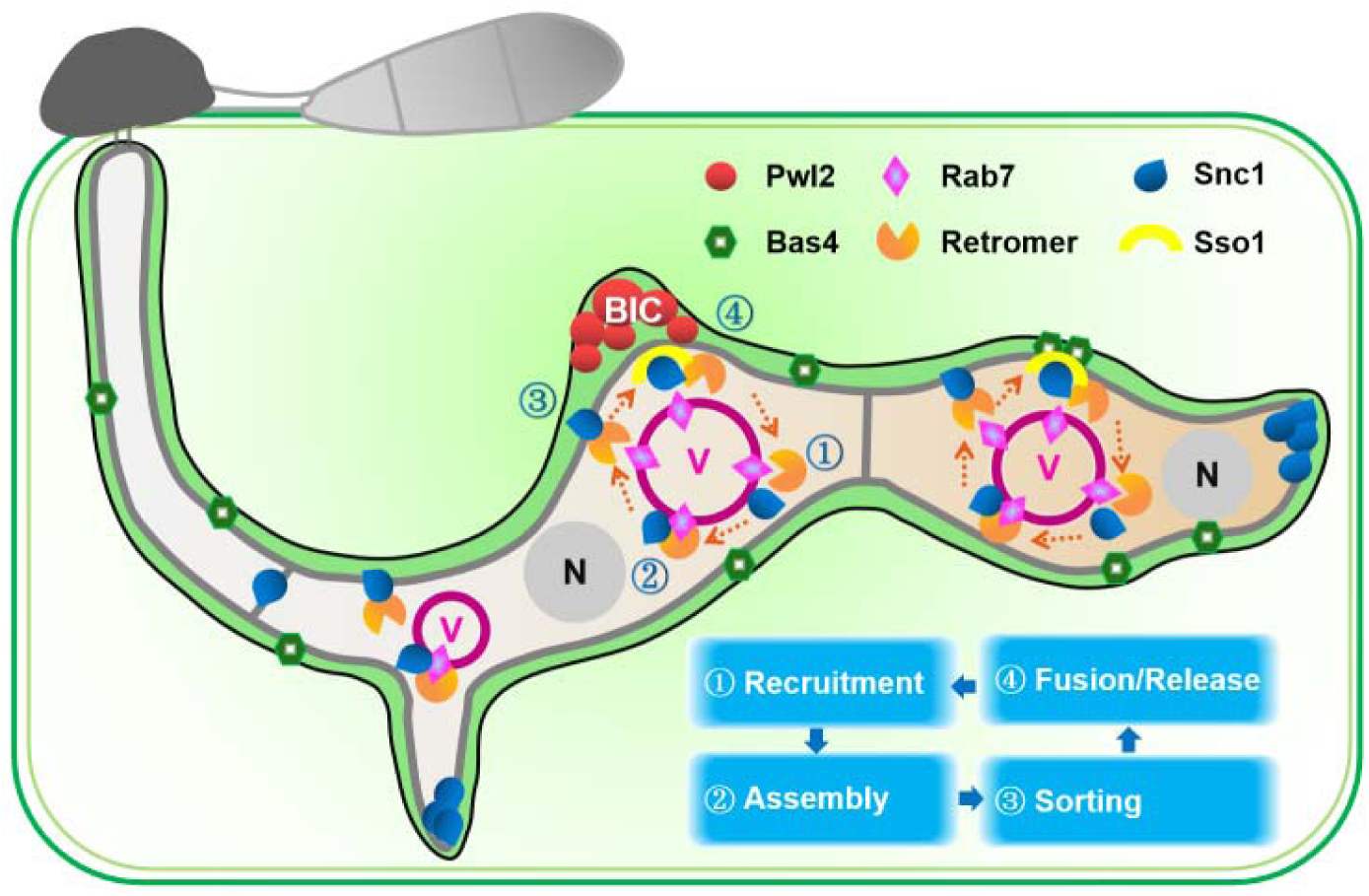
A proposed model depicting the effector secretion in MoRab7/Retromer/MoSnc1-based endolysosomal cascade. During *M. oryzae* biotrophic invasion, cytoplasmic effectors (eg. Plw2) preferentially accumulate in the biotrophic interfacial complex (BIC), while it is often associated with a large vacuole (purple) in bulbous invasive hyphae, compared to nucleus (N). MoRab7 uniformly localizes to vacuolar membrane and recruits retromer complex for vesicular trafficking. A conserved v-SNARE protein, MoSnc1, is recognized by retromer complex and subjected to recycling from vacuole to BIC associated membrane and fungal plasma membrane, mediating fusion of secretory vesicles with this target membrane. The correct/efficient secretion of cytoplasmic and apoplastic effectors (eg. Bas4) require MoRab7/Retromer/MoSnc1-based endolysosomal cascade.

Vacuole (or lysosome) is the central organelle responsible for degradation, and nutrient signaling, which plays important roles in cellular homeostasis. Vacuole receives various cargo proteins from different pathways and further decides the destiny of the cargoes to either degrade or to recycle [28, 39, 40]. Live cell imaging helped establish that a prominent vacuole is the most adjacent organelle to the BIC structure throughout *M. oryzae* invasion and colonization (Fig. 1). Since the MoRab7 GTPase is the major regulator of vacuole biogenesis and the homotypic fusion machinery [28, 41], we further investigated the association between MoRab7 and the BIC development. Our results showed that MoRab7 stays in close proximity to the BIC and its dominant negative variant significantly impairs the interface leading to defects in plant infection (Fig. 2). Although the vacuole is adjacent to the BIC, we failed to observe a direct connection between the vacuole *per se* and the BIC structure. Since the vacuole has been regarded as a signal transduction center [40], we hypothesized that a trafficking machinery is likely recruited to the vacuole and is responsible for transportation between vacuole and BIC. Interestingly, the retromer complex was further identified to be such trafficking machinery between vacuole and the BIC. Retromer is normally required for retrograde trafficking of cargoes from endosomes to the Golgi, and recent studies have highlighted its role in mediating trafficking to plasma membrane too [19, 20]. However, such retromer function in the phytopathogenic fungi has remained unexplored thus far. We provide evidence that the retromer vesicles are able to attach to the plasma membrane and BIC-associated regions during invasive growth. In addition, retromer is responsible for the trafficking and membrane-targeting of the v-SNARE MoSnc1, which is required for BIC development and effector secretion (Fig. 7, 8 and 9). We provide evidence that vacuole is associated with the BIC structure and the MoRab7-retromer-MoSnc1 cascade is responsible for the trafficking events between the vacuole and biotrophy interface. To date, the knowledge about the relationship between vacuole and BIC have remained limited. Whether the vacuole serves as a storage hub or a regulation/recycling center for BIC development and effector secretion needs further investigation. Additionally, super-resolution live cell microscopy techniques are needed to further define the details of the vacuole-BIC association; and to define the origin and destination(s) of vesicular traffic and to differentiate various membranes at and near the biotrophy interface.

SNARE (soluble N-ethylmaleimide-sensitive factor attachment protein receptor) proteins are considered as the engine for membrane fusion events essential for various cellular processes [42]. Accumulating evidence has indicated the important roles of SNAREs in fungal development and pathogenicity of the rice blast fungus. MoSec22, an R-SNARE, was shown to be involved in cell wall integrity, conidiogenesis and pathogenicity [43]. The vacuolar SNARE MoVam7 plays important roles in regulating vacuole formation and membrane trafficking needed for fungal development and virulence [44]. The t-SNARE MoSso1 is involved in the BIC structure development and cytoplasmic effector secretion [5]. In addition, the syntaxin Syn8 is required for intracellular transport and secretion of avirulence effectors Avr-Pia and Avr-Pizt. However, how the SNARE proteins are regulated and the role of other SNAREs in fungal development and pathogenicity in *M. oryzae* remains unclear. Through pull down assays using MoVps35-GFP, we identified 12 SNARE proteins using mass spectrometry (Table1, Table S1), thus underlying a potential relationship between the retromer and SNARE proteins. Among these SNARE proteins, MoSnc1 was further characterized and the role of MoSnc1 with retromer and its function in BIC structure development and effector secretion were uncovered. Consistent with a previous study, MoSnc1 localizes to hyphal tip, septum, plasma membrane and subapical BIC-associated cells (Fig. 8 and Fig. S12, [5]), while it was missorted to vacuole in *ΔMovps35* mutant. In addition, MoVps35 physically interacts with MoSnc1 in both vegetative and invasive growth periods (Fig. 7). These results indicated the important role of the retromer complex in intracellular trafficking of MoSnc1. Since MoSnc1 displays BIC-associated localization, we further charactered the role of MoSnc1 in effector secretion. Interestingly, MoSnc1 plays dual functions in secretion of apoplastic and cytoplasmic effectors. Similar to Δ*Mosso1* mutant [5], loss of MoSnc1 also disrupts the BIC structure development (Fig. 8 and Fig. 9). Thus, we shed new light on the regulatory mechanisms and the role of v-SNARE MoSnc1 in proper and timely effector secretion in *M. oryzae*. However, knowledge about the SNARE function in phytopathogens remains far from complete, and future expansion in such research will certainly further our understanding of how SNAREs function in pathogenic development and in fungus-host interaction.

Generally, proteins with signal peptides are secreted through the conventional ER-to-Golgi pathway enroute the plasma membrane. While the leaderless cargoes (proteins without signal peptide) are secreted via the unconventional pathway [8, 13]. Autophagy is an evolutionarily conserved process required for protein degradation and recent studies have expanded its role in unconventional protein secretion [45–47]. In *M. oryzae*, autophagy is thought to be involved in programed cell death, nutrient homeostasis, and stress adaption [48, 49]. However, the relationship between autophagy and the unconventional protein secretion remains largely unexplored. The sorting nexins, Rab GTPases and SNARE proteins play key roles in autophagy [50–54], and are also required for pathogenicity in *M. oryzae*. The sorting nexin Snx41 is required for glutathione-based antioxidant defense and pexophagy, and loss of Snx41 significantly reduces *Magnaporthe* virulence [55, 56]. MoSec4, a small Rab GTPase, is required for proper localization of the cytoplasmic effector Pwl2 [57]. MoRab7 and retromer are both required for autophagy [23, 30], and we provide evidence that MoRab7-retromer-MoSnc1 cascade is essential for effector secretion in the rice blast fungus (Fig. 10). According to previous reports, various autophagy interactors play important roles in effector secretion, including vacuole protein Imp1, the Sec61β, endocytosis related protein MoEnd3 and the putative verprolin protein MoVrp1 [58–60]. It is possible that the autophagy process is associated with effector secretion in *M. oryzae*, and understanding of such fascinating association would shed new insights on the complex membrane trafficking events needed for precise effector secretion in phytopathogenic fungi.

## Materials and Methods

### Fungal strains, medium, and growth conditions

All *Magnaporthe oryzae* strains used in this study are listed in the Supporting Information Table S2. The *M. oryzae* strains were stored as dried filter paper stocks at −20°C, and cultured on complete medium (CM) and/or prune agar (PA) at 25°C under continuous light for 10 days as described previously [61].

### DNA manipulation and fungal transformation

*M. oryzae* transformants are described in Supplemental Table S2. Details about plasmid construction are listed in Supplemental Table S3, and oligonucleotide primers provided in Supplemental Table S4. All fusion constructs were verified by DNA sequencing. Plasmids of interest were transferred to the *M. oryzae* laboratory strain using Agrobacterium tumefaciens-mediated transformation or PEG-mediated protoplast transformation [62]. For generation of *MoSNC1* gene deletion mutants, about 1-kb upstream and downstream sequences of *MoSNC1* were amplified using specific primers (see Supplemental Table S4). We then generated the *MoSNC1* gene replacement constructs by using split-marker approach [23].

### Staining procedures and cytological analyses

For staining of the vacuolar lumen, CMAC (7-amino-4-chloromethylcoumarin, Cat. #C2110, Invitrogen) was used at a final concentration of 10μM. FM4-64 (Cat. #T3166, Invitrogen) was used at a final concentration of 10 μM for staining the Spitzenkörper, plasma membrane, septa as well as the endosomes. To examine cytological dynamics during *M. oryzae-*rice interaction, rice sheaths (cv. CO39) were inoculated with conidia from *M. oryzae* strains expressing epifluorescently-labeled effectors and/or the other indicated proteins (Table S2). Rice leaf sheath inoculation with *M. oryzae* was performed as described previously [63, 64]. In brief, leaf sheaths were detached from 4-week-old rice seedlings, and were cut into 7-8 cm segments. Inoculation was performed by injecting *M. oryzae* conidia (5 × 10^4^ conidial/mL) in inner interior of rice leaf sheaths. The inoculated sheaths were placed on a 96-well PCR plate and were incubated under high humidity (95%) and at 25°C for about 20-48 hour. For microscopic inspection, the inoculated leaf sheaths were trimmed into ultrathin sections and imaged using the spinning disk confocal microscope, or Nikon A1 laser scanning confocal microscope (Nikon, Japan). The spinning-disk confocal microscope was equipped with a Yokogawa CUS-X1 spinning-disk confocal system and a 100×/1.4Lnumerical aperture (NA) oil lens objective. The images were captured using a 16-bit digital Orca-Flash4.0 scientific complementary metal oxide semiconductor (sCMOS) camera (Hamamatsu Photonics KK). Image processing was performed using Imaris (Bitplane) and Fiji (https://imagej.net/Fiji). The excitation/emission wavelengths used were 405 nm/425-475nm for CMAC, 488 nm/500–550 nm for GFP, and 561 nm/570–620 nm for mCherry and FM4-64.

### Affinity purification and mass spectrometric analysis

For affinity purification, mycelia of the wild type Guy11, Guy11-GFP and MoVps35-GFP were harvested and ground into fine powder in liquid nitrogen. The related samples were lysed in extracted buffer (10 mM Tris HCl pH 7.5, 150 mM NaCl, 0.5 mM EDTA, 1% Triton X100, 2 mM PMSF) containing Protease Inhibitor Cocktail (Sangon Biotech, Shanghai, CN). Total proteins were inoculated with 30 μL GFP-Trap®_A beads (ChromoTek Inc., Hauppauge, NY, USA) at 4°C for 4h and then the bound proteins eluted by heating at 100°C for 10 min. The mass spectrometric analysis was carried as previously described [38].

### Immunoblot and co-immunoprecipitation (Co-IP) assays

For collection of vegetative hyphae, mycelial plugs of the indicated strains were cultured in CM liquid medium at 28°C, 110 rpm for 3 days. For collection of invasive hyphae, hyphal blocks of the indicated strains were inoculated on 7-day-old barley leaves for 2 days. Briefly, the indicated samples were ground into fine powder and then lysed in extraction buffer containing Protease Inhibitor Cocktail (Sangon Biotech, Shanghai, CN). Total proteins were then incubated with 30 μL GFP-Trap®_A beads (ChromoTek Inc., Hauppauge, NY, USA) at 4°C for 4h. Similarity, the bound proteins were eluted by adding protein loading buffer and then heating at 100°C for 10 min. The resulting protein samples were separated by 10% SDS-PAGE and then analyzed by western blotting with anti-GFP antibody (GFP-Tag (7G9) Mouse mAb, Abmart, Shanghai, CN), anti-Flag antibody (DYKDDDDK-Tag(3B9) mAb, Abmart, Shanghai, CN), Goat Anti-Mouse IgG HRP (Abmart, Shanghai, CN) and Anti-Myc antibody (HRP Anti-Myc tag antibody, Abcam, Cambridge, MA, USA).

### Tet-Off gene expression system

The Tet-Off system was engineered by fusing the promoter and coding region of MoVps35 to pFGL1252_TetGFP(Hyg) vector (Addgene ID 118992), which contains a specific Tet-off cassette activated by tetracycline or doxycycline. Briefly, the promoter sequence of MoVps35 was amplified using primer MoVps35 Tet-AF and MoVps35 Tet-AR, and the resulting fragment ligated to pFGL1252_TetGFP(Hyg) vector (digested with *Xho* I and *Eco*R I) to generated the pFGL1252_TetGFP(Hyg)-Vps35A construct. Next, ∼1500 bp coding region of MoVps35 was amplified using primers MoVps35 Tet-BF and MoVps35 Tet-BR and then ligated into pFGL1252_TetGFP(Hyg)-Vps35A (digested with *Pst* I and *Hin*d III) to generate the pFGL1252_TetGFP(Hyg)-Vps35AB construct. The resulting construct was transformed into *M. oryzae* wild type protoplasts and the transformants were verified by PCR and sequencing in addition to the presence of the requisite epifluorescence. The correct/positive transformants were further verified by confocal microscopy and phenotype analysis with tetracycline or doxycycline treatment.

### Phenotypic analysis of mutants

For growth assays, the indicated strains were cultured in CM, PA and SYM medium at 28°C for 10 days. Conidiation and conidiophore formation assays were performed on rice bran agar plates at 28°C with periodic exposure to 12-h light/dark cycles. To induce the formation of appressoria, conidia of the indicated strains were inoculated on hydrophobic coverslips for 8 to 24h. For plant infection assays, hyphal block or spore suspension (1 × 10^5^ conidia/mL) of the indicated strains was inoculated on intact or wounded barley leaves for 5 days. For spray assays, 5 mL conidial suspension of the indicated strains (1×10^5^ conidia/mL) were inoculated on 3-week-old rice for 7 days. For invasive growth assays, spores of the indicated strains were inoculated on rice leaves sheaths at 26°C for 36h and then observed under laser confocal microscopy.

## Acknowledgements

This work was supported by the National Science Fund for Excellent Young Scholars (32122071), the National Natural Science Foundation of China (31772106), and the Natural Science Foundation of Fujian Province (2021J06015). Research in the Naqvi lab is supported by grants from Temasek Life Sciences Laboratory and the National Research Foundation, Singapore. All authors declared no financial or other potential conflict of interest.

---

**Figure S1.**
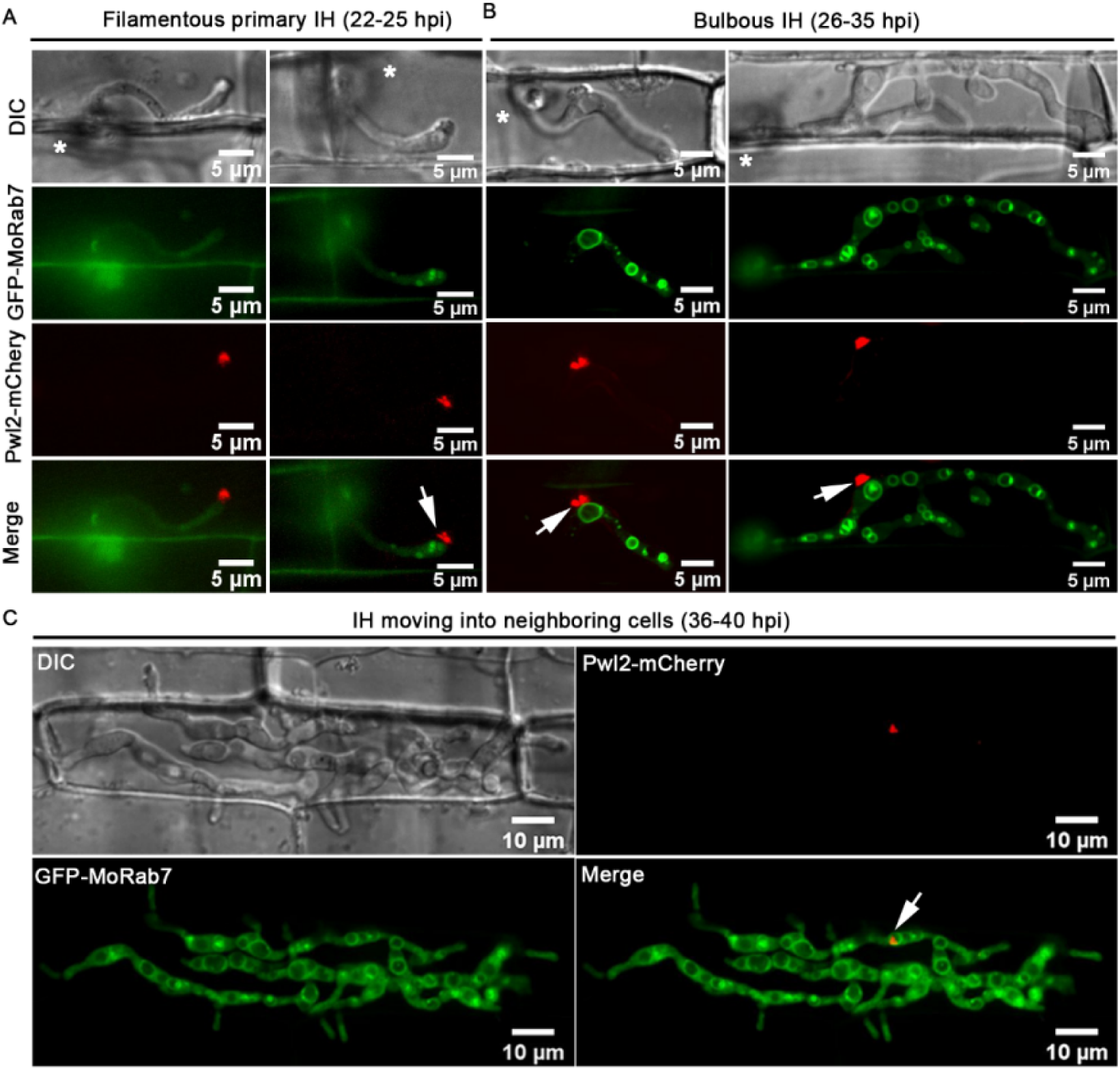
Close association of MoRab7 and BIC during host invasion in the blast fungus. (A) Localization of GFP-MoRab7 and Pwl2-mCherry at the early infection stage (22-25 hpi). At the initial penetration stage, Pwl2-mCherry is positioned at the tip of the primary filamentous invasive hyphae (IH) while GFP-MoRab7 signal is quite weak. Once the tip of the primary hypha begins to swell, the GFP-MoRab7 signal intensifies and enhances, and gradually emerges. (B) At the mid infection stage (26-35 hpi), the GFP-MoRab7 positive ring structure is adjacent to the BIC (arrow) in the bulbous hyphae. (C) At the later infection stage (36-40 hpi), GFP-MoRab7 ring structure associates with the BIC (arrow) as the IH grow into the neighboring cells. Asterisks indicate appressoria.

**Figure S2.**
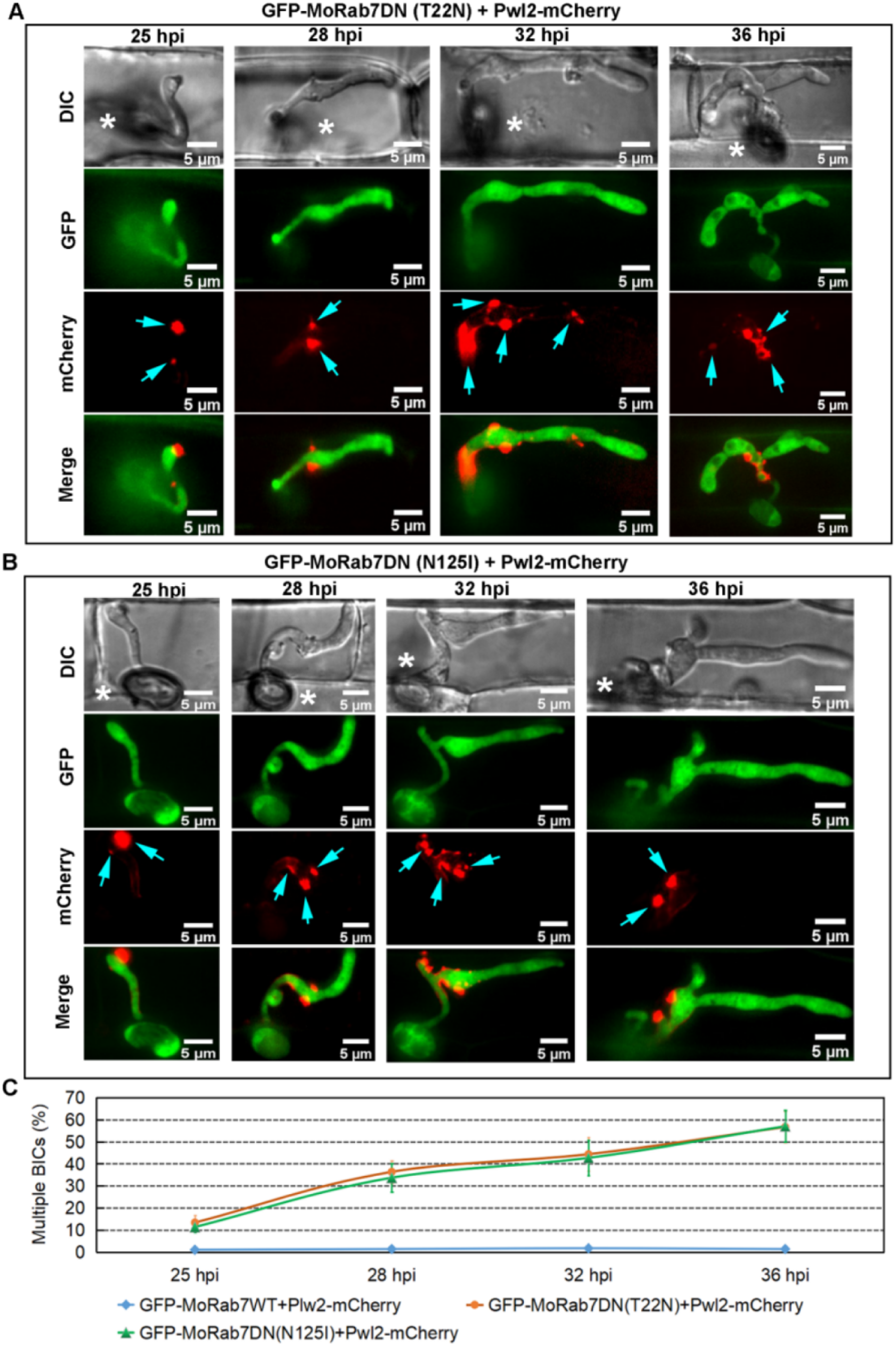
Inactivation of MoRab7 impairs Pwl2 localization. (A) The dominant-negative GFP-MoRab7^T22N^ mutation (locked in GDP-bound state) using *PWL2* promoter failed to target the MoRab7 protein to the vacuolar membrane; it rather caused inappropriate secretion of the protein. Multiple and discrete fluorescent foci (arrows) are observed in the IH, especially in the bulbous IH. (B) Another dominant-negative site (N125I) of MoRab7 was selected and overexpressed using PWL2 promoter. The result shows that GFP-MoRab7DN(N125I) are present at the cytosol and cause improper secretion of Pwl2-mCherry protein in the bulbous IH. Asterisks indicate appressoria. (C) A line graph showing the percentage of multiple BICs of the IH in the indicated strains at each time point. Error bars represent standard deviations; three biological replicates; 300 infected cells observed.

**Figure S3.**
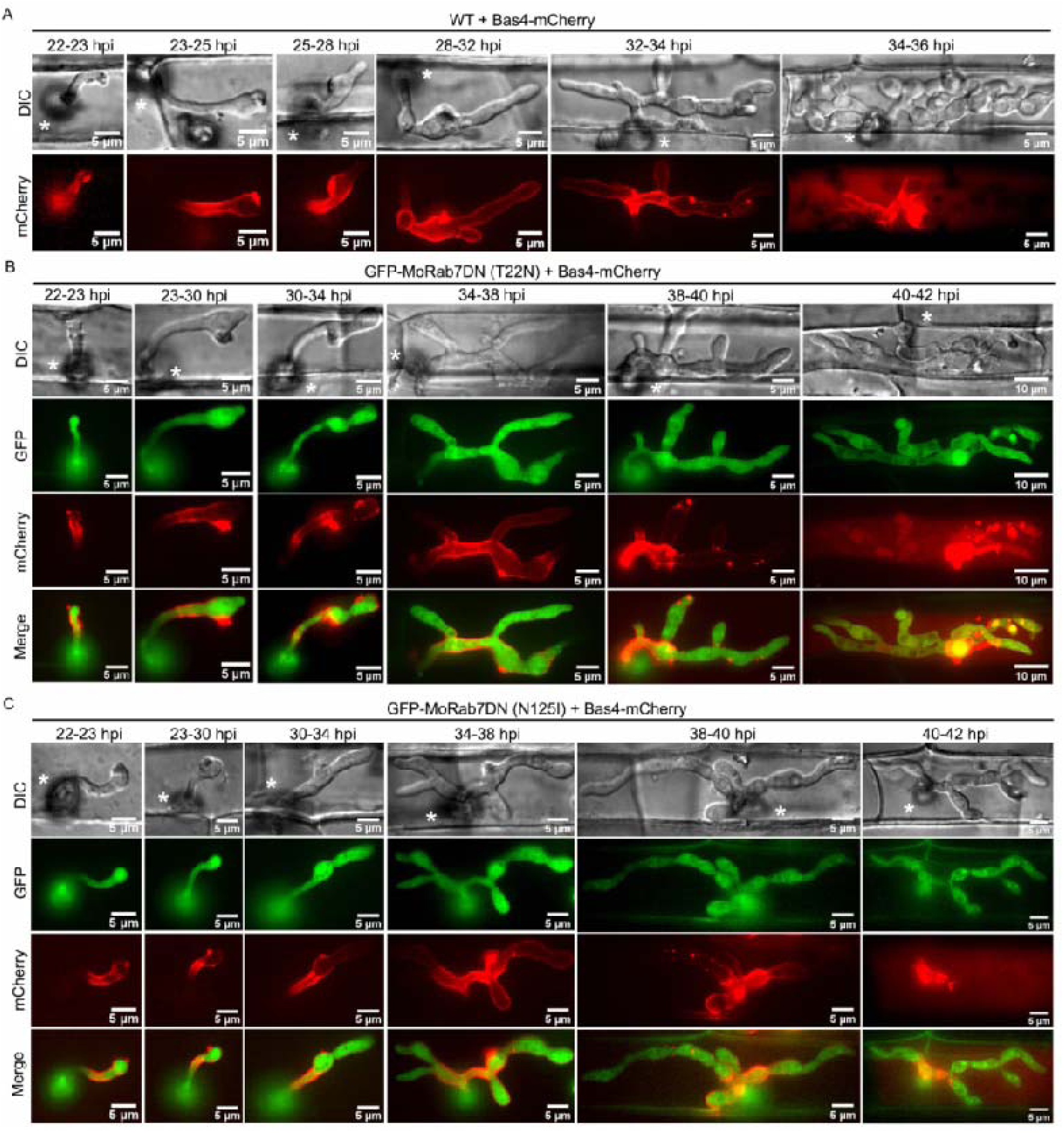
Inhibition of MoRab7 GTPase activity does not affect the localization of the apoplastic effector Bas4 during invasive growth in rice sheath. (A) The Bas4-mCherry fusion protein was expressed under its native promoter in the wild type (WT) and initially outlines the invasive hyphae during invasion of the initial epidermal cells (22-32 hpi). After 32 h, Bas4-mCherry signals appear patchy or diffused inside the infected rice cells, suggesting that EIHM lost its integrity. (B and C) Overexpression of dominant-negative GFP-MoRab7 via T22N or N125I (locked in GDP-bound state) mutation using PWL2 promoter resulted in failure of the protein to target vacuolar membrane and in delay of IH development, but does not affect the normal secretion of Bas4-mCherry protein. Asterisks indicate appressoria.

**Figure S4.**
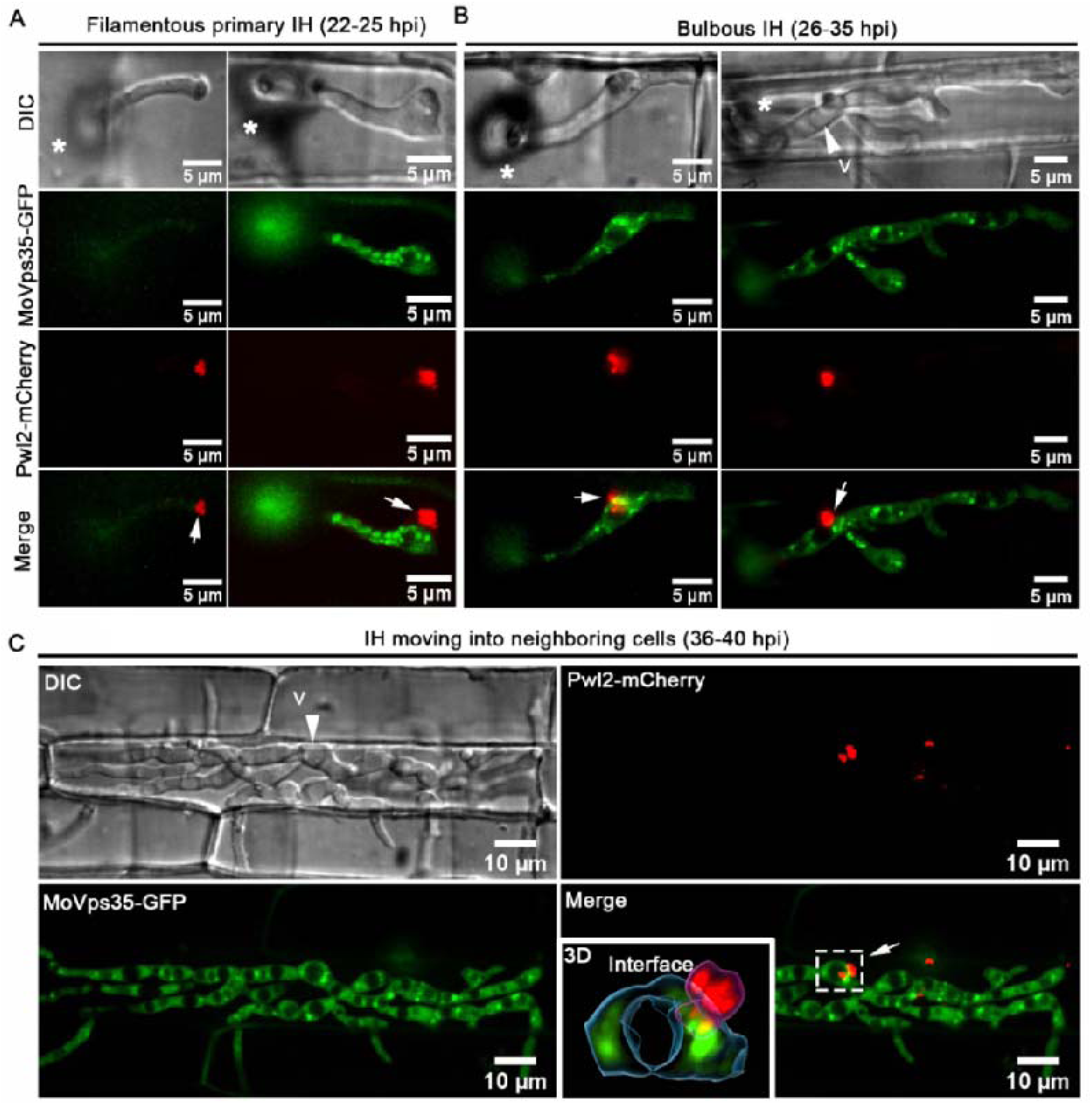
MoVps35 shows punctate epifluorescence in close proximity to the BIC. (A) At the initial penetration stage (22-25 hpi), Pwl2-mCherry is concentrated at the tip BIC preceding the MoVps35 expression in filamentous invasive hyphae. As the primary filamentous IH begin to swell, the MoVps35-GFP showed distinct fluorescence and some of which are near to the BIC. (B) At the mid penetration time points (26-35 hpi), filamentous primary IH have developed into bulbous IH and MoVps35 localizes to the vacuolar membrane next to the BIC. (C) At the later infection stage (36-40 hpi), MoVps35 is adjacent to the vacuolar membrane and also associates with the BIC (arrow) as the IH grow into neighboring cells. The 3D image shows an intimate interfacial positioning of the vesicular sorting machinery (green vesicles, MoVps35-GFP) and the BIC (red vesicles, Pwl2-mCherry). V indicates vacuole (vacuoles appear hollow in DIC images). Asterisks indicate appressoria while the white arrows point to the BIC.

**Figure S5.**
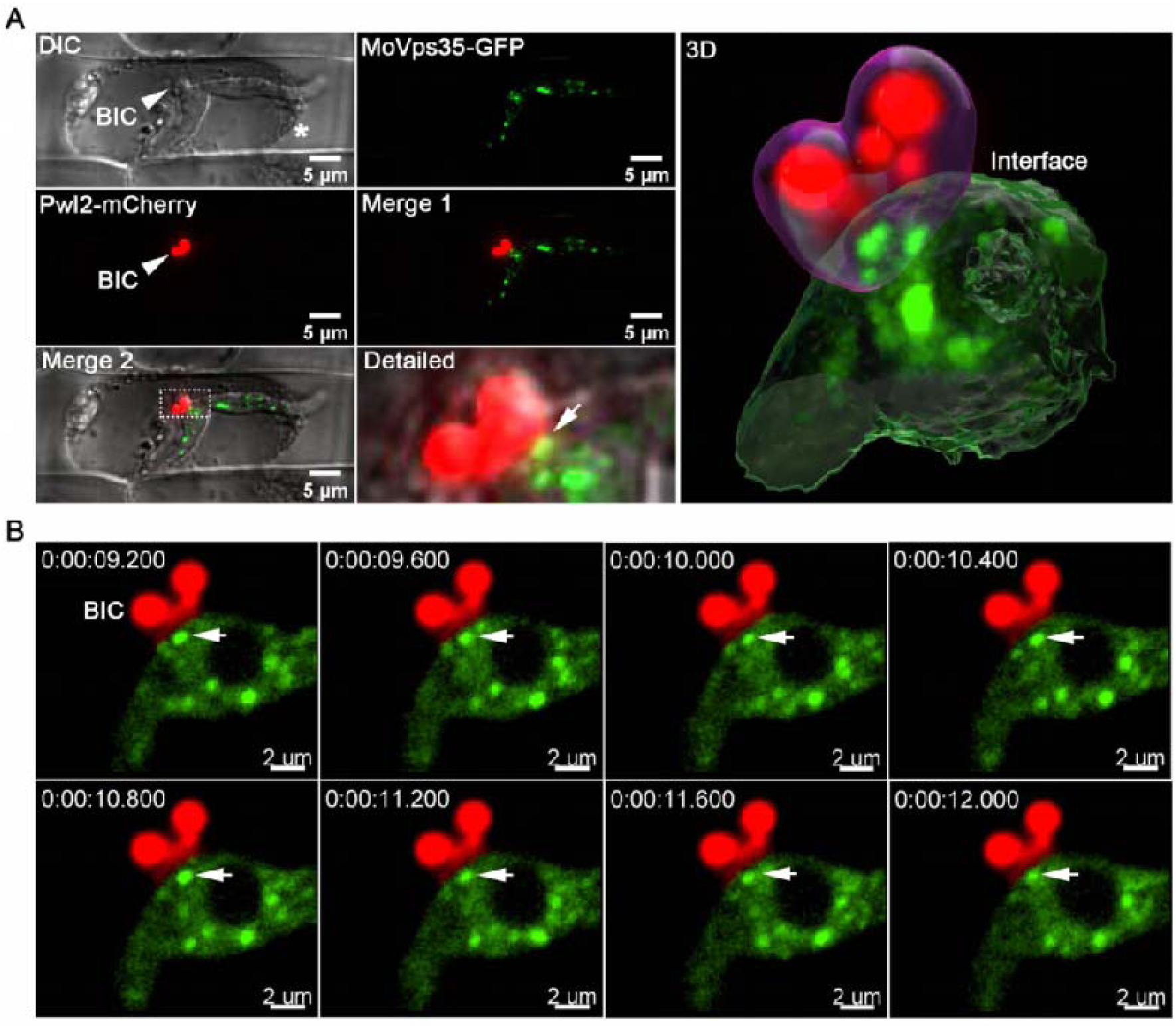
MoVps35-GFP-labeled vesicles localize proximal to the BIC upon sucrose-induced plasmolysis. (A) Visualization of bulbous IH expressing MoVps35-GFP and Pwl2-mCherry in the plasmolyzed rice cell. The surface rendered 3D image (panel on the right) shows the close positioning of the vesicular sorting machinery (green vesicles, MoVps35-GFP) and the BIC (Pwl2-mCherry). (B) Dynamic tracking of MoVps35-GFP labeled vesicles near the BIC during host invasion (Supplementary video 13). Images were captured at 400-ms intervals. Asterisk indicates shrunken protoplast from the rice cell; arrowheads point to the BIC; and full arrows to the MoVps35-GFP labeled vesicles.

**Figure S6.**
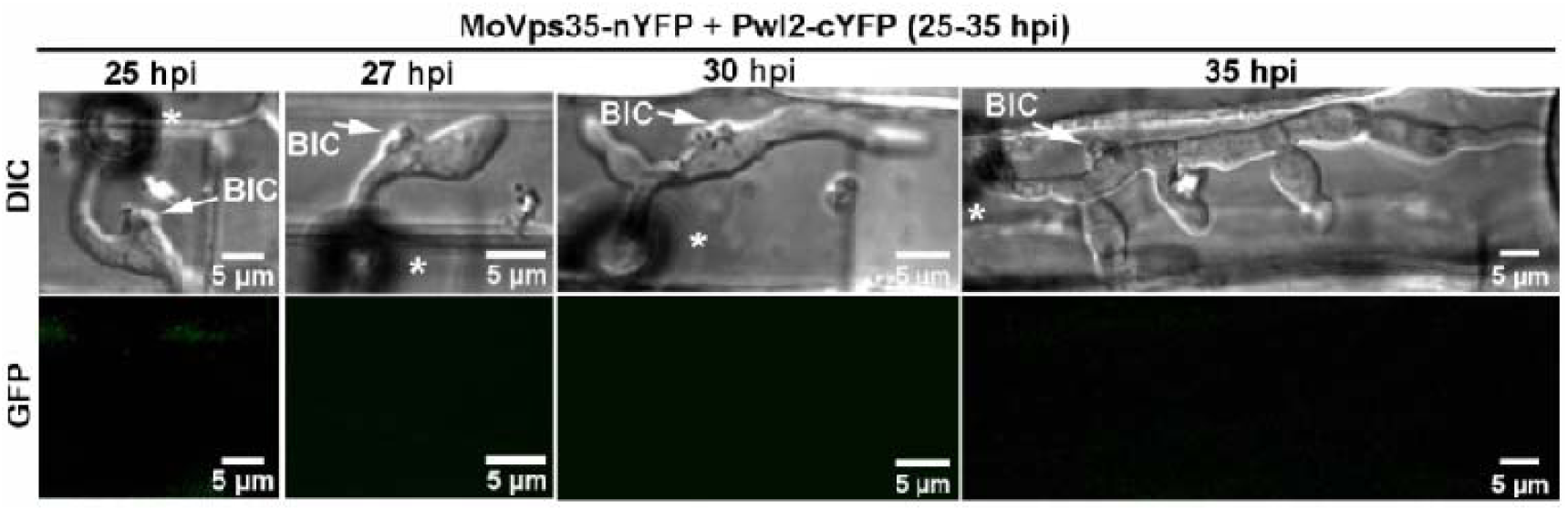
Bimolecular fluorescence complementation (BiFC) assay to test the interaction between MoVps35 and Pwl2 *in vivo*. No obvious fluorescent signal was observed in the IH co-expressing MoVps35-nYFP and Pwl2-cYFP during *M. oryzae* infection (26-35 hpi). Arrows point to the BIC; asterisks indicate appressoria. DIC: Differential Interference Contrast images.

**Figure S7.**
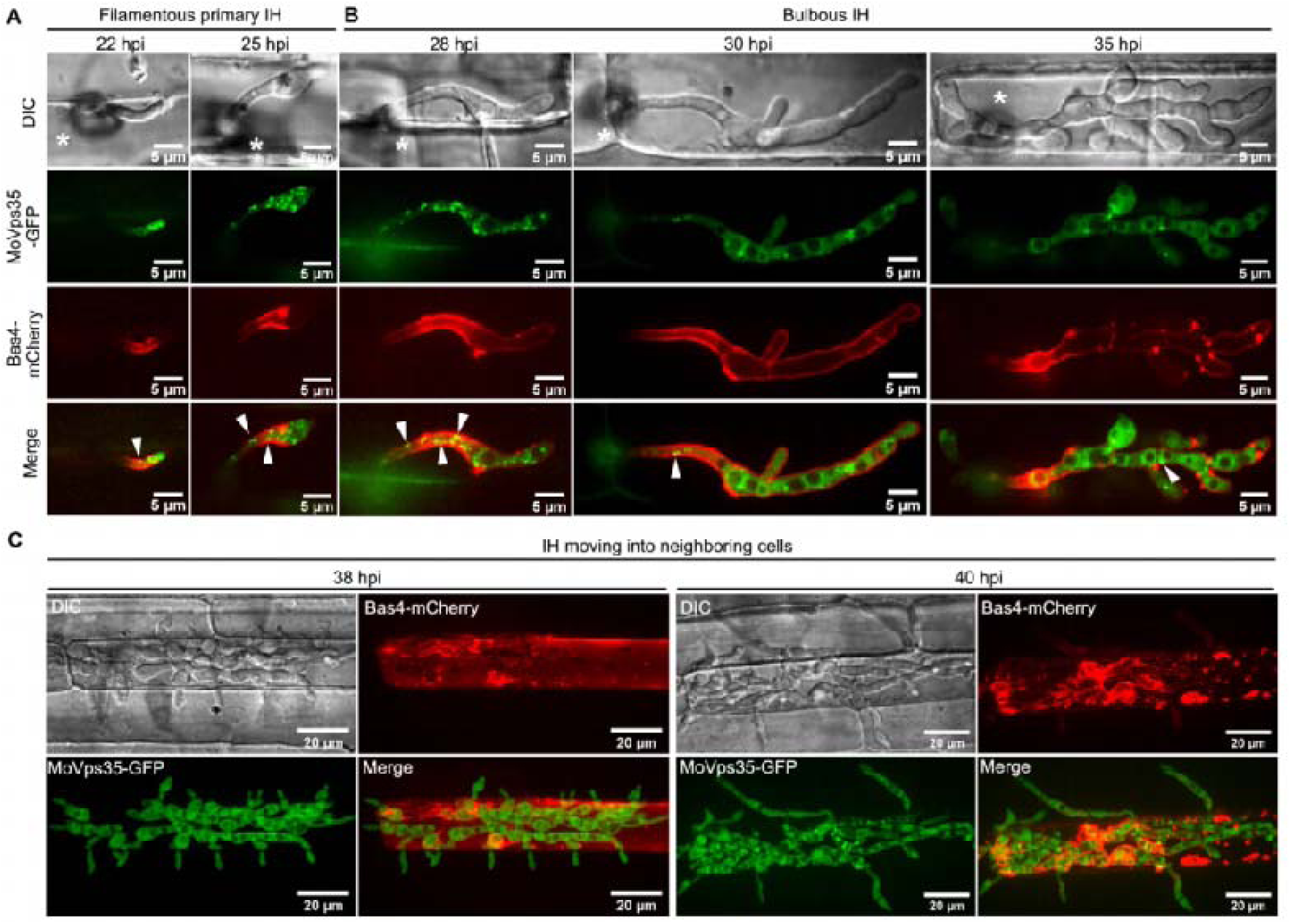
Visualization of the infectious hyphae co-expressing MoVps35-GFP and Bas4-mCherry (EIHM-specific marker) during *M. oryzae* infection. (A and B) Some of MoVps35-GFP-labeled dots are adjacent to the EIHM (arrowheads) during invasion of the initial rice sheath cells (22-35 hpi). (C) At later time points (>38 hpi), the IH entered the neighboring cells, and Bas4-mCherry is evident in the plant cytoplasm but does not diffuse into the adjacent rice cells after the EIHM collapses. Asterisks indicate appressoria.

**Figure S8.**
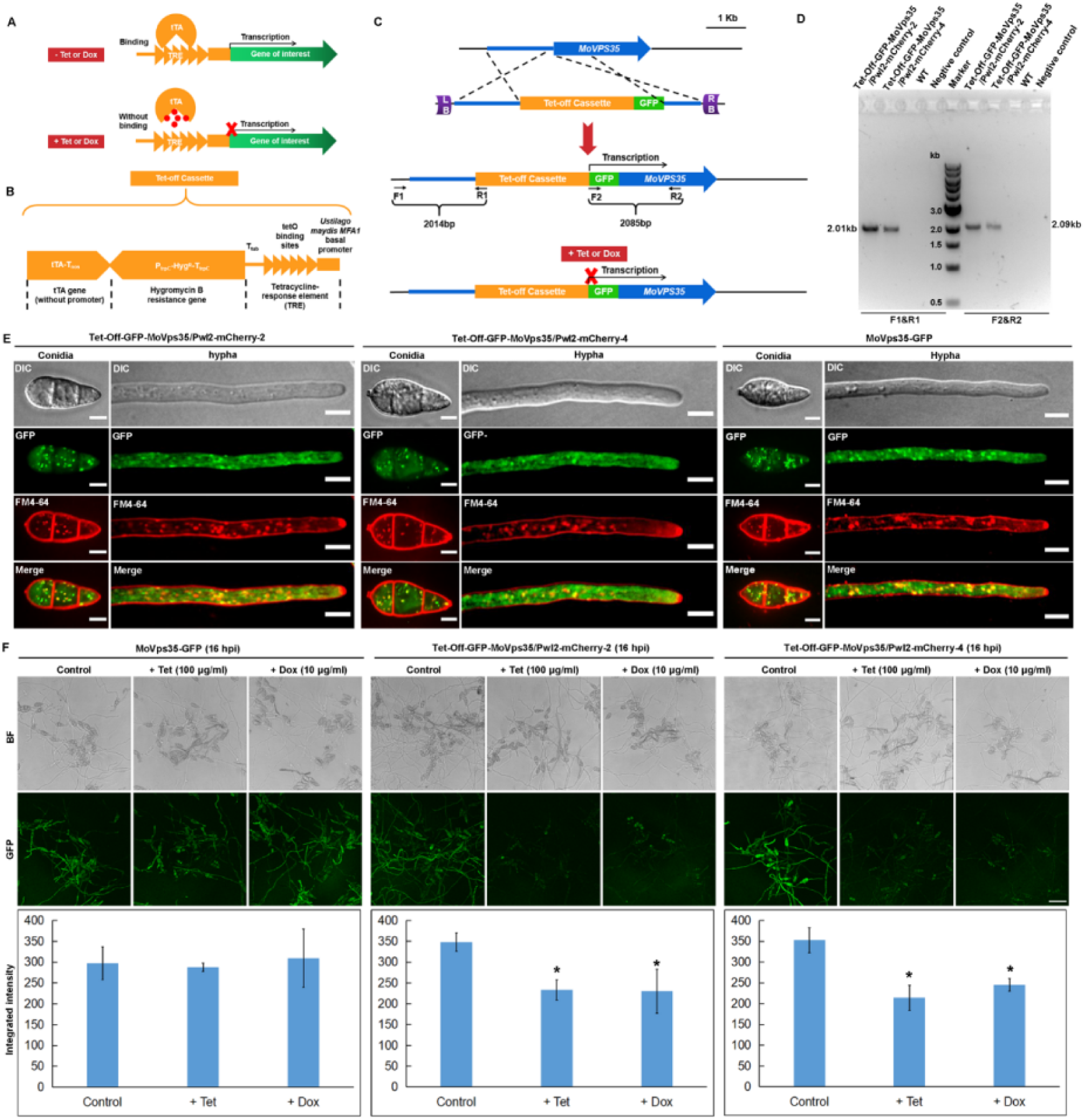
Generation and verification of Tet-off-GFP-MoVps35/Pwl2-mCherry strains. (A) Schematic representation of the gene regulon of the Tet-off system. In the absence of Tet (tetracycline) or Dox (doxycycline), tTA (tetracycline-dependent transactivator) binds the TRE (Tet-response element) and activates gene transcription. (B) Schematic representation of Tet-Off cassette consisting of the tetracycline-dependent transactivator-encoding gene tTA (without promoter) and the tetracycline-responsive element TRE separated by the Hygromycin B-resistance locus. (C) Schematic representation of MoVps35-locus integration via homologous recombination. Primers F1, R1, F2 and R2 were used to screen and identify the correct transformants. (D) PCR-based verification of the Tet-off-GFP-MoVps35/Pwl2-mCherry strains. DNA bands of 2.01-kb and 2.09-kb were observed in the two independent Tet-off-GFP-MoVps35/Pwl2-mCherry strains, respectively, while no bands were observed in the wild-type (WT) strain or the negative control. (E) Analysis of the localization of MoVps35 from two independent Tet-off-GFP-MoVps35/Pwl2-mCherry strains. GFP-MoVps35 localizes to punctate structures that partially co-localized with FM4-64 that marks the endosomal membranes in conidia and hyphae, which is consistent with the localization of MoVps35-GFP in the complemented strain. (F) Tetracycline- or doxycycline-regulated *MoVPS35* expression. Treatment of Tet-off-GFP-MoVps35/Pwl2-mCherry strain with either tetracycline or doxycycline (100 or 10 µg/ml, respectively) significantly reduced the GFP-MoVps35 expression to basal levels. In the control strain MoVps35-GFP, neither the presence of tetracycline nor that of doxycycline influenced the GFP intensity.

**Figure S9.**
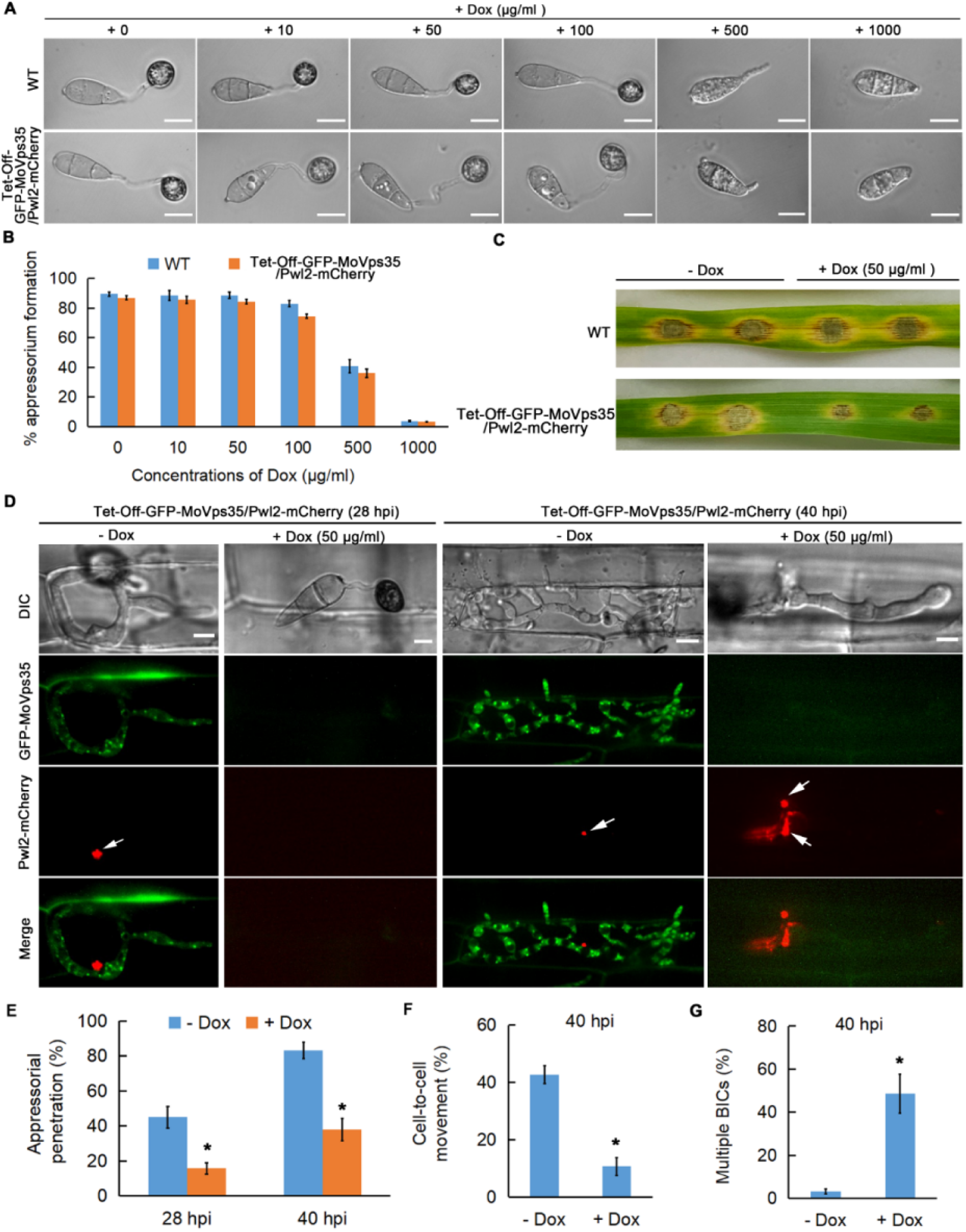
Retromer-dependent regulation of plant infection and Pwl2 effector secretion by *M. oryzae.* (A and B) Addition of Dox (doxycycline; <100 µg/ml) does not affect appressorium development (Student’s *t* test; three biological replicates; 600 spores observed). (C) Leaf drop inoculation assay showing the effect of Dox (50 µg/ml) on lesion formation by the Tet-off-GFP-MoVps35/Pwl2-mCherry strain. The presence of Dox has no significant effect on the wild type. (D) Chemico-genetic inactivation of MoVps35 impairs plant infection and Pwl2 effector secretion in *M. oryzae.* Representative laser confocal micrographs showing strong GFP-MoVps35 expression at 28 and 40 hpi in IH during infection. The GFP-MoVps35 expression ceased after treatment with 50 μg/ml Dox. Consistently, addition of Dox to the conidia of Tet-off-GFP-MoVps35/Pwl2-mCherry strain results in a significant decrease in appressorium-mediated host penetration (28 hpi) and increased the overall number of BICs in the invasive hypha (40 hpi). BICs are indicated by arrows. (E) A bar chart showing the frequency of plant penetration at 28 and 40 hpi by *M. oryzae* conidia in the presence or absence of 50 μg/ml Dox (*P < 0.05; Student’s *t* test; three biological replicates; 300 appressorium observed). (F) Bar chart depicting the frequency of cell-to-cell movement at 40 hpi by *M. oryzae* in the presence or absence of 50 μg/ml Dox (*P < 0.05; Student’s *t* test; three biological replicates; 150 infected cells observed). (G) A bar chart showing the frequency of multiple BICs at 40 hpi in *M. oryzae* in the presence or absence of 50 μg/ml Dox (*P < 0.05; Student’s *t* test; three biological replicates; 150 infected cells observed).

**Figure S10.**
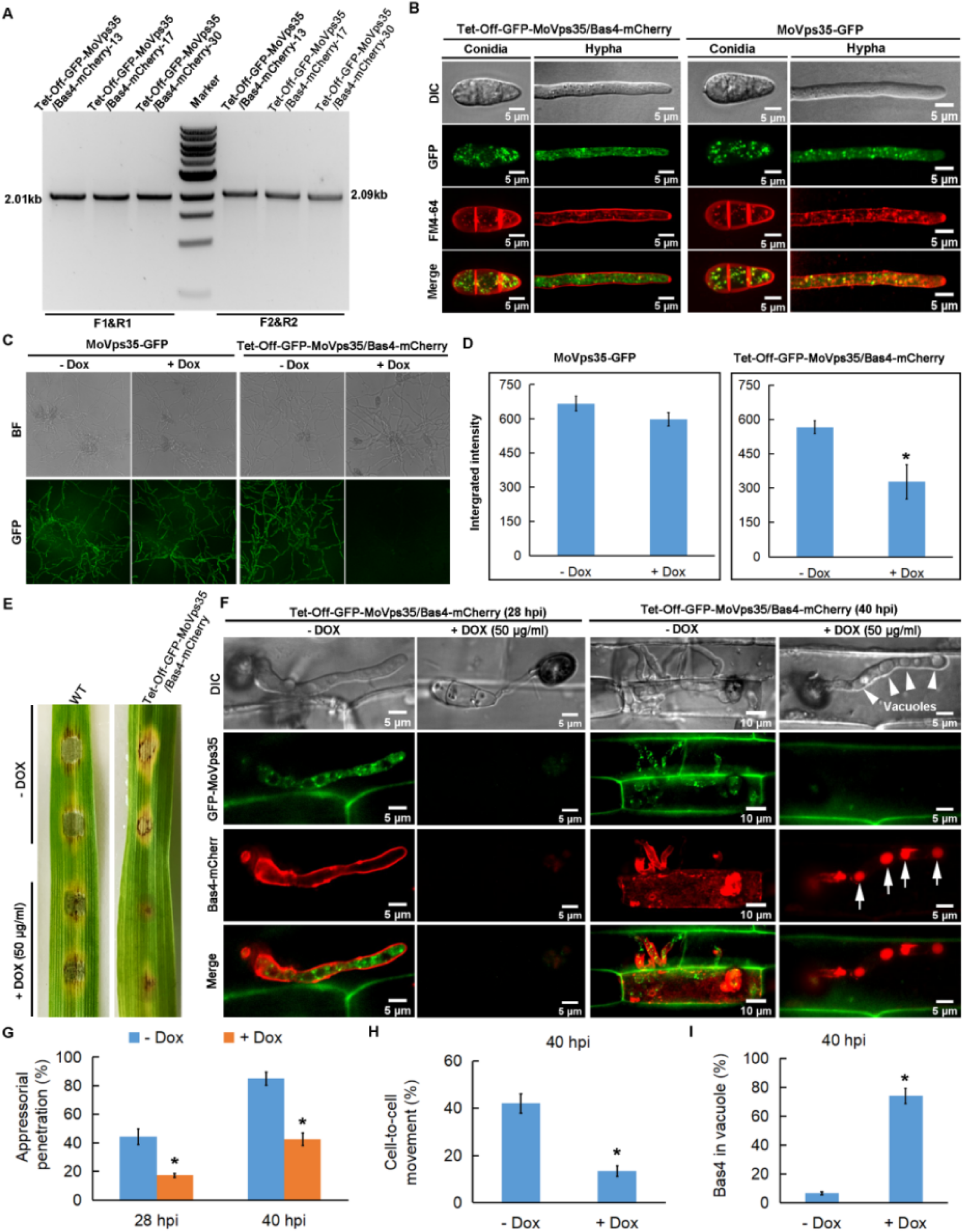
Generation of Tet-off-GFP-MoVps35/Bas4-mCherry strain and testing its role in plant infection and Bas4 secretion. (A) PCR verification of the Tet-off-GFP-MoVps35/Bas4-mCherry positive transformants. DNA bands of 2.01-kb and 2.09-kb were observed in the three independent Tet-off-GFP-MoVps35/Bas4-mCherry strains, respectively. (B) Analysis of the localization of MoVps35 in Tet-off-GFP-MoVps35/Bas4-mCherry strain. The GFP-MoVps35 punctae partially colocalize with FM4-64 that marks the endosomal membranes in conidia and hyphae, consistent with the localization of MoVps35-GFP in the complemented strain. (C) Doxycycline-regulated MoVps35 expression. Treatment of Tet-off-GFP-MoVps35/Bas4-mCherry strain with doxycycline (10 µg/ml) reduced the GFP expression to basal levels. In the control MoVps35-GFP strain, treatment with doxycycline did not influence the GFP intensity. (D) Quantification of GFP intensity. (E) Leaf drop inoculation assay showing the effect of Dox (50 µg/ml) addition on lesion formation by the Tet-off-GFP-MoVps35/Bas4-mCherry strain. The presence of Dox has no significant effect on the WT. (F) Chemical genetic inactivation of MoVps35 impairs plant infection and Bas4 effector secretion by *M. oryzae.* Representative laser confocal micrographs showing strong GFP-MoVps35 expression at 28 and 40 hpi in IH undergoing infection. The GFP-MoVps35 expression ceased after treatment with 50 μg/ml Dox. Consistently, addition of Dox to the conidia of Tet-off-GFP-MoVps35/Bas4-mCherry resulted in a significant decrease in appressorium function of host penetration (28 hpi) and caused mislocalization of Bas4 into the vacuoles in *M. oryzae* (40 hpi). Arrowheads point to the vacuoles (vacuoles appear as hollows in DIC imaging). Mislocalized Bas4 effector is indicated by arrows. (G) A bar chart showing the frequency of plant penetration at 28 and 40 hpi by *M. oryzae* conidia in the presence or absence of 50 μg/ml Dox (*P < 0.05; Student’s *t* test; three biological replicates; 300 appressorium observed). (H) Bar chart showing the frequency of cell-to-cell movement at 40 hpi by *M. oryzae* in the presence or absence of 50 μg/ml Dox (*P < 0.05; Student’s *t* test; three biological replicates; 150 infected cells observed). (I) A bar chart showing the percentage of *M. oryzae* IH containing vacuolar Bas4-mCherry at 40 hpi in the presence or absence of 50 μg/ml Dox (*P < 0.05; Student’s *t* test; three biological replicates; 150 infected cells observed).

**Figure S11.**
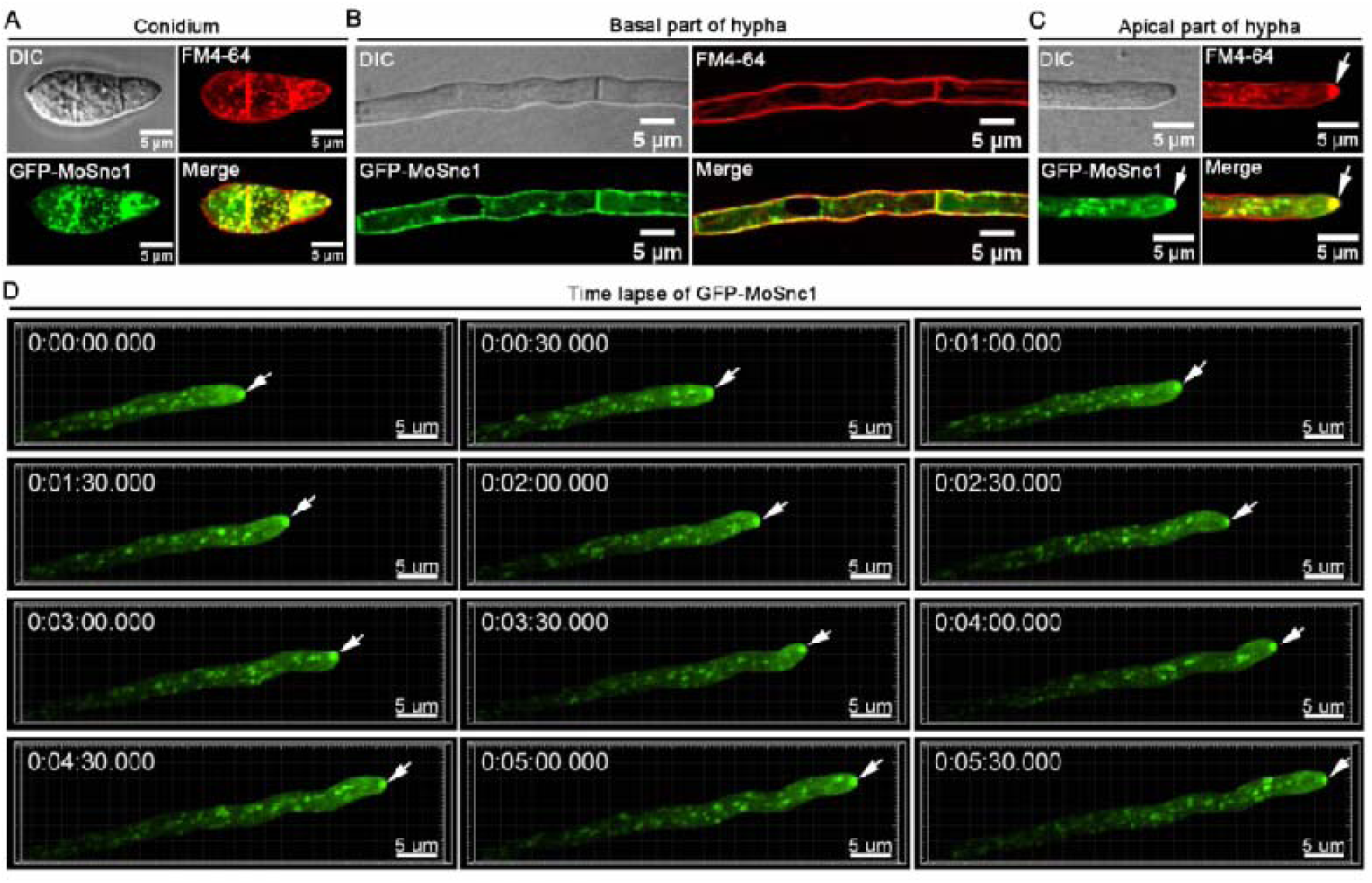
GFP-MoSnc1 localizes to endosomes, plasma membrane and hyphal apex. (A) GFP-MoSnc1 signal was found in multiple punctae inside a conidium and were partially co-localized with the endocytic and endosome marker dye FM4-64. (B) In basal part of hyphae, GFP-MoSnc1 is mainly localized to the plasma membrane and septa stained with FM4-64. (C) In the growing hypha, GFP-MoSnc1 signal also concentrated at the hyphal apex and is co-localized with the FM4-64 marked Spitzenkörper and/or polarisome (arrow). (D) Dynamic tracking of GFP-MoSnc1 in the vegetative hyphae growing on CM medium (Supplementary video 16). DIC: Differential Interference Contrast.

**Figure S12.**
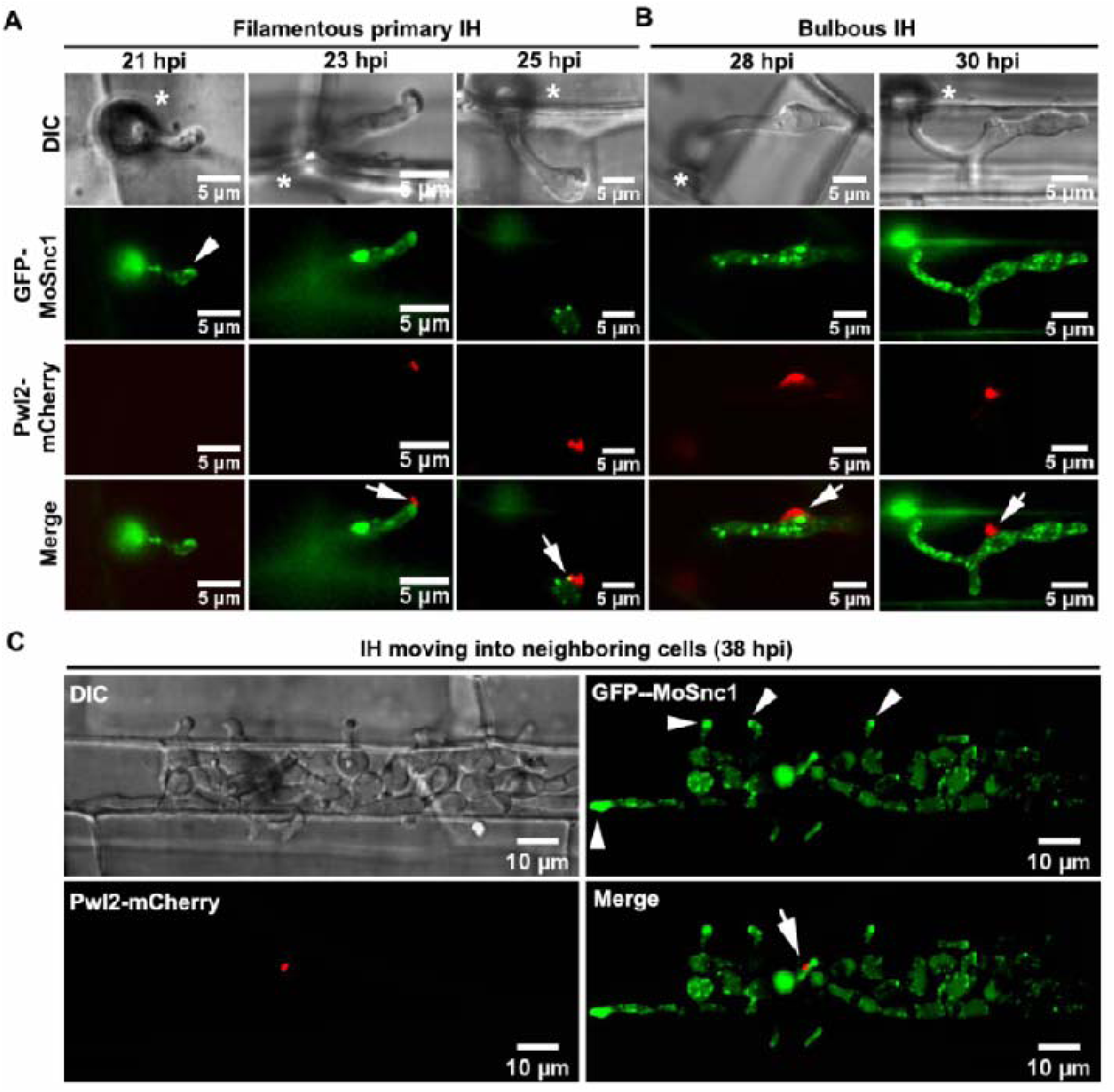
MoSnc1 shows punctate fluorescence next to the BIC. (A) At the initial penetration stage (21 hpi), GFP-MoSnc1 signal is evident at the tip (arrowhead) of the filamentous invasive hyphae while Pwl-mCherry is undetectable at this time point. When primary invasive hyphae begin to swell, the GFP-MoSnc1 fluorescence is still concentrated at the hyphal tip where the BIC (arrow) is eventually formed (23 hpi). As the hypha develops, the BIC is repositioned to the side of the first IH cell, while GFP-MoSnc1 still associates with the BIC (arrow) (25 hpi). (B) At 26-30 hpi, the filamentous primary IH develop into bulbous IH where some of the GFP-MoSnc1 vesicles are closely associated with the BIC (arrow). (C) At the later infection stage (38 hpi), GFP-MoSnc1 vesicles remain associate with the BIC. In addition, GFP-MoSnc1 is concentrated at the IH apex (arrowhead) as the hyphae grow into the neighboring cells. V indicates the vacuole. Asterisks indicate the appressoria. Arrows mark the BIC. Arrowheads indicate the IH apex/tip.

**Figure S13.**
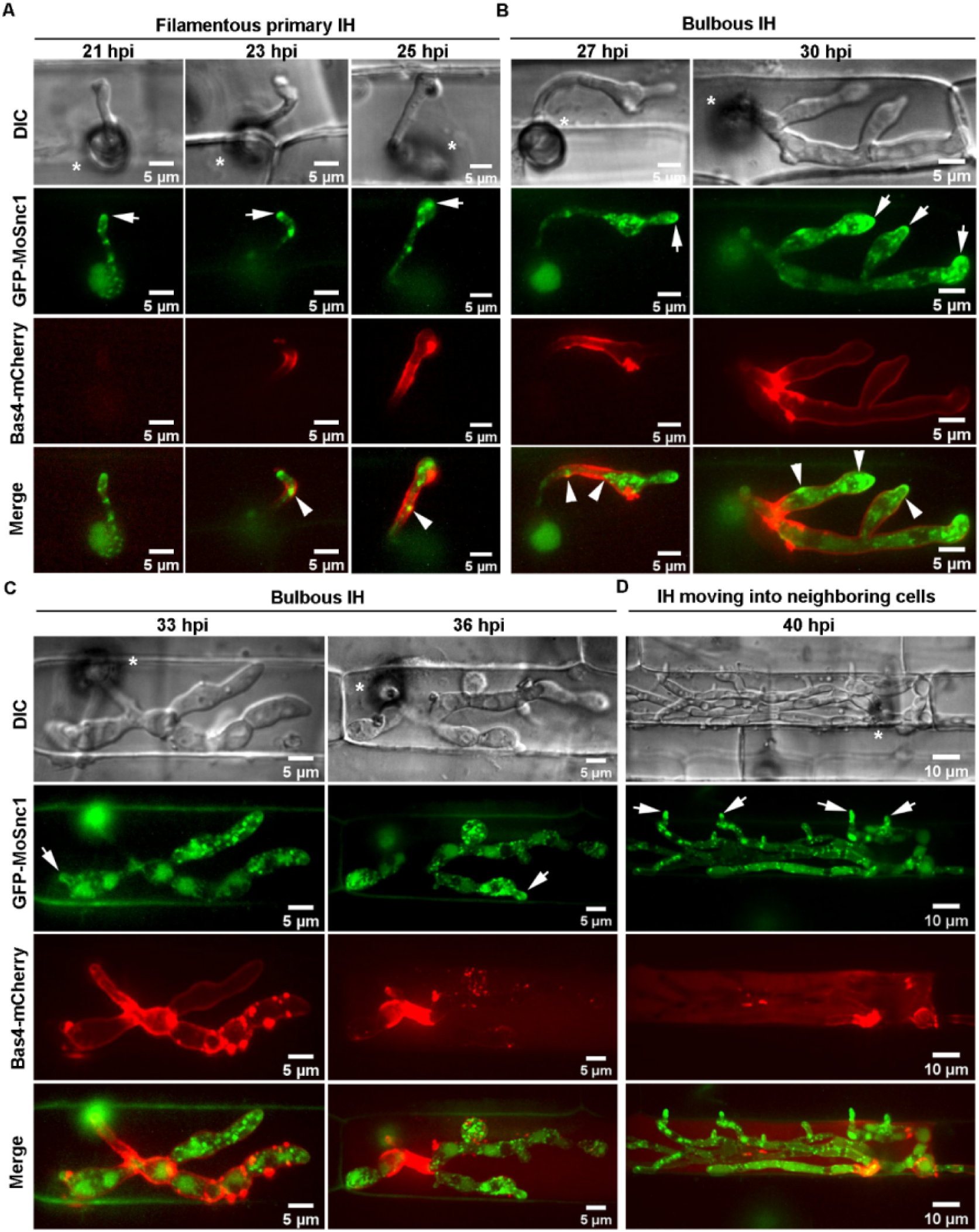
Visualization of infectious hyphae expressing GFP-MoSnc1 and Bas4-mCherry during host infection by *M. oryzae*. (A and B) Confocal micrographs showing some of the GFP-MoSnc1-labeled dots positioned adjacent to the EIHM (arrowheads) during invasion of the initial rice sheath cells (21-30 hpi). (C) Bas4-mCherry signals appear patchy at 33 hpi and further diffuse inside the infected rice cells at 36 hpi, suggesting loss of EIHM integrity. (D) At the later time point (40 hpi), when the IH spread to the neighboring cells, Bas4-mCherry fluorescence is observed in the host cytoplasm but does not diffuse into the adjacent rice cells after the EIHM collapses. Arrows indicate accumulation of GFP-MoSnc1 at the tip of the invasive hyphae. Asterisks indicate appressoria.

**Figure S14.**
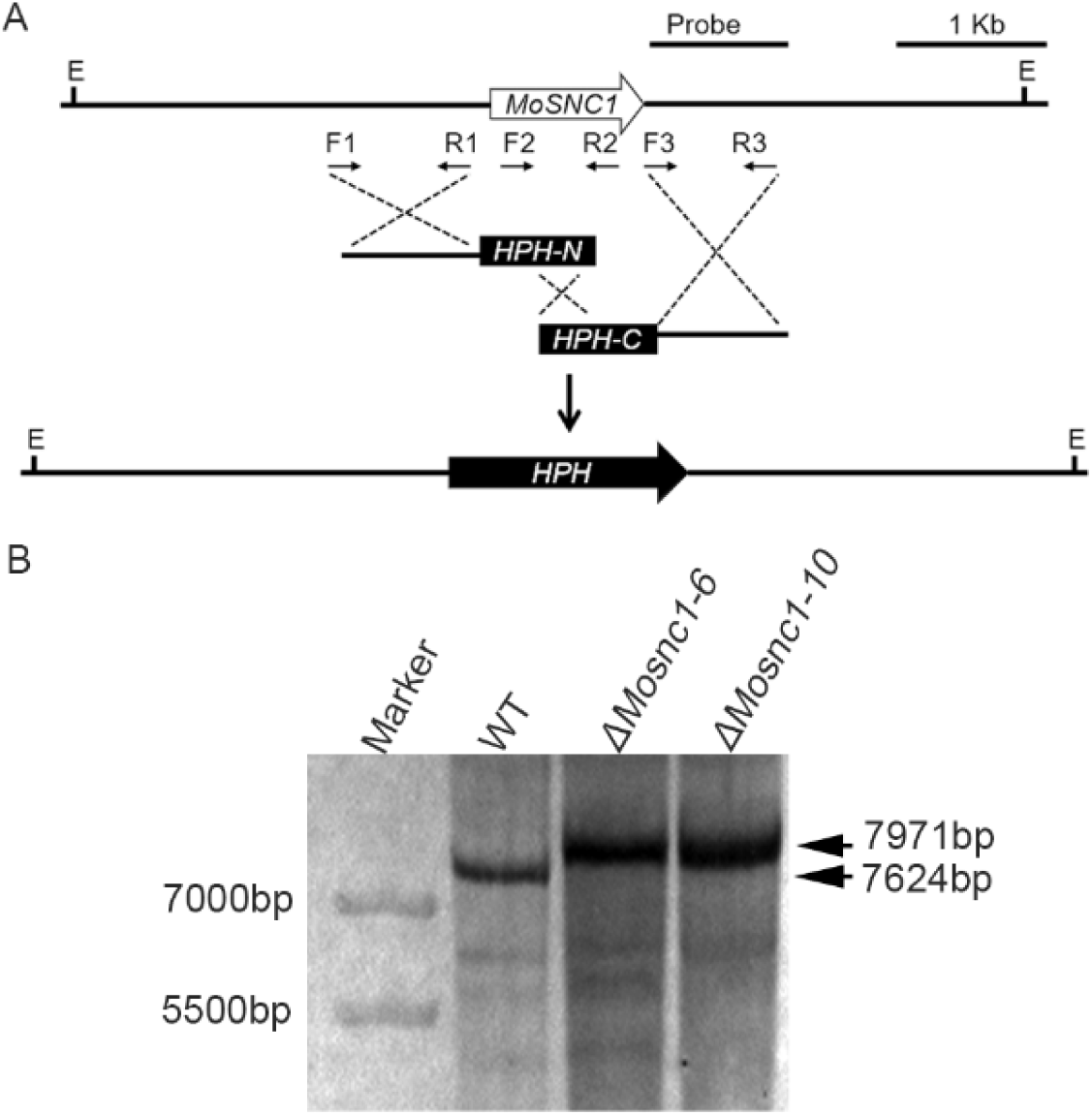
Schematic diagram of the genomic regions of *MoSNC1* and the *HPH* (Hygromycin B phosphotransferase) locus. Primers F1, R1, F3 and R3 were used to generate the *MoSNC1* gene replacement construct, and F2 and R2 were used for mutant screening and identification. E, *Eco*RI. (B) Southern blots of *Eco*RI-digested genomic DNA were probed with the 3’UTR fragment of *MoSNC1*. A 7624bp band was observed in the wild-type, while 7971bp band was observed in the two independent mutants.

**Figure S15.**
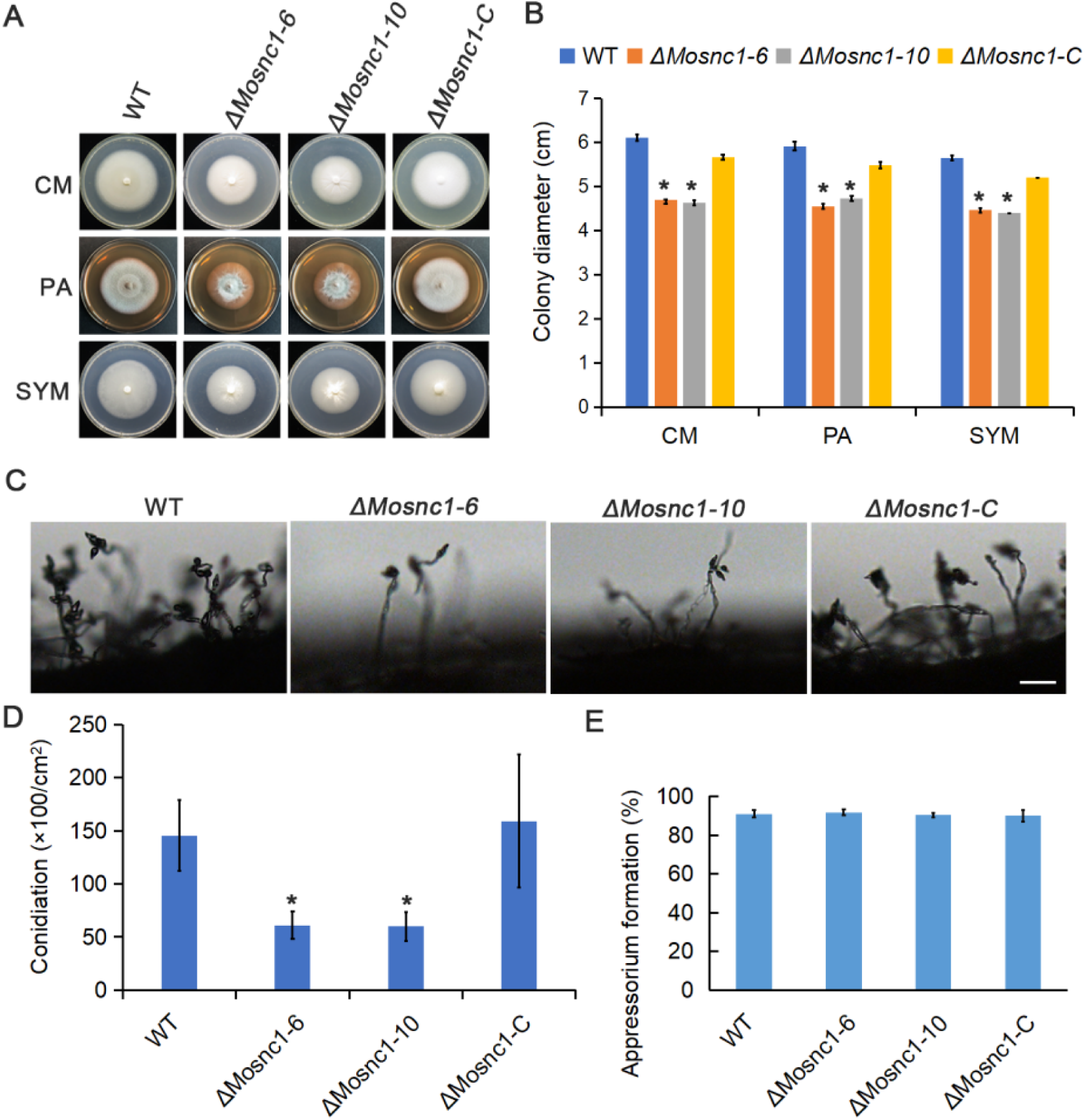
Functional analyses of *ΔMosnc1* mutants. The *ΔMosnc1* mutants displayed reduced mycelial growth on CM, PA and SYM media. (B) Statistical analysis of mycelial growth after incubation of the indicated strains on CM, PA and SYM media. Data represent mean ± SD from three independent replicates, and asterisks indicate statistically significant differences (P<0.05). (C) The number of conidia on conidiophores is reduced in the *ΔMosnc1* mutant. Bar = 50 μm. (D) Quantification of conidia. The respective strains were initially grown in the dark for three days followed by exposure to constant illumination for 14 day on PA plate (diameter 9 cm). Data represent mean ± SD from three independent replicates, and asterisk indicate statistically significant differences (P<0.05). (E) Quantification of appressorium formation. Data represent mean ± SD from three independent replicates.

Video S1. 3D carousel view showing the positioning of BIC, vacuole and nucleus during host invasion.

Video S2. Dynamics of GFP-MoRab7- and Pwl2-mCherry-marked BIC at the early stage of host invasion.

Video S3. Dynamics of GFP-MoRab7- and Pwl2-mCherry-marked BIC at the mid stage of host invasion.

Video S4. 3D carousel view of infectious hyphae expressing GFP-MoRab7WT and Pwl2-mCherry

Video S5. 3D carousel view of infectious hyphae expressing GFP-MoRab7DN(T22N) and Pwl2-mCherry.

Video S6. 3D carousel view of infectious hyphae expressing GFP-MoRab7DN(N125I) and Pwl2-mCherry.

Video S7. 3D carousel view of infectious hyphae expressing GFP-MoRab7CA(Q67L) and Pwl2-mCherry.

Video S8. Dynamics of mCherry-MoRab7 and MoVps35-GFP during rice cell invasion by *M. oryzae*.

Video S9. A 3D carousel view showing the intimate interfacial positioning of the vesicular sorting machinery (green vesicles, MoVps35-GFP) and the BIC (red vesicles, Pwl2-mCherry) at the early stage of host invasion.

Video S10. A 3D carousel view showing the intimate interfacial positioning of the vesicular sorting machinery (green vesicles, MoVps35-GFP) and the BIC (red vesicles, Pwl2-mCherry) at the mid stage of host invasion.

Video S11. Dynamic tracking of the retromer complex subunit MoVps35 transport to the BIC-associated zone during host invasion.

Video S12. Dynamic tracking of the retromer complex subunit MoVps17 transport to the BIC-associated zone during host invasion.

Video S13. Dynamic tracking of MoVps35-GFP-labeled vesicles transports to the BIC-adjacent region after sucrose-induced plasmolysis.

Video S14. 3D carousel view of infectious hyphae expressing MoVps35-GFP and Bas4-mCherry.

Video S15. Dynamic tracking of the retromer complex subunit MoVps35-GFP transport to the EIHMx-associated zone during host invasion.

Video S16. Polarized localization of GFP-MoSnc1 during hyphal extension.

Video S17. Dynamics of GFP-MoSnc1- and Pwl2-mCherry-marked BIC at the early stage of host invasion.

Video S18. Dynamic tracking of the GFP-MoSnc1 transport to the BIC-associated zone during host invasion.

Video S19. A 3D carousel view confirms that GFP-MoSnc1-labeled vesicles are covered by the BIC during host invasion.

Video S20. 3D carousel view the indicated box showing close positioning of GFP-MoSnc1- and Bas4-mCherry-marked EIHMx.

Video S21. 3D carousel view of the indicated box showing close positioning of GFP-MoSnc1- and Bas4-mCherry-marked EIHMx.

## Notes

### Competing Interest Statement

The authors have declared no competing interest.

## References

1. Dean R, Van Kan JA, Pretorius ZA, Hammond-Kosack KE, Di Pietro A, Spanu PD, et al. The Top 10 fungal pathogens in molecular plant pathology. Mol Plant Pathol. 2012;13(4):414–30. Epub 2012/04/05. doi: 10.1111/j.1364-3703.2011.00783.x. PubMed PMID: 22471698; PubMed Central PMCID: PMCPMC6638784.

2. Giraldo MC, Valent B. Filamentous plant pathogen effectors in action. Nat Rev Microbiol. 2013;11(11):800–14. Epub 2013/10/17. doi: 10.1038/nrmicro3119. PubMed PMID: 24129511.

3. Tariqjaveed M, Mateen A, Wang S, Qiu S, Zheng X, Zhang J, et al. Versatile effectors of phytopathogenic fungi target host immunity. J Integr Plant Biol. 2021;63(11):1856–73. Epub 2021/08/13. doi: 10.1111/jipb.13162. PubMed PMID: 34383388.

4. Fernandez J, Orth K. Rise of a Cereal Killer: The Biology of Magnaporthe oryzae Biotrophic Growth. Trends Microbiol. 2018;26(7):582–97. Epub 2018/02/06. doi: 10.1016/j.tim.2017.12.007. PubMed PMID: 29395728; PubMed Central PMCID: PMCPMC6003838.

5. Giraldo MC, Dagdas YF, Gupta YK, Mentlak TA, Yi M, Martinez-Rocha AL, et al. Two distinct secretion systems facilitate tissue invasion by the rice blast fungus Magnaporthe oryzae. Nat Commun. 2013;4:1996. Epub 2013/06/19. doi: 10.1038/ncomms2996. PubMed PMID: 23774898; PubMed Central PMCID: PMCPMC3709508.

6. Mentlak TA, Kombrink A, Shinya T, Ryder LS, Otomo I, Saitoh H, et al. Effector-mediated suppression of chitin-triggered immunity by magnaporthe oryzae is necessary for rice blast disease. Plant Cell. 2012;24(1):322–35. Epub 2012/01/24. doi: 10.1105/tpc.111.092957. PubMed PMID: 22267486; PubMed Central PMCID: PMCPMC3289562.

7. Mosquera G, Giraldo MC, Khang CH, Coughlan S, Valent B. Interaction transcriptome analysis identifies Magnaporthe oryzae BAS1-4 as Biotrophy-associated secreted proteins in rice blast disease. Plant Cell. 2009;21(4):1273–90. Epub 2009/04/10. doi: 10.1105/tpc.107.055228. PubMed PMID: 19357089; PubMed Central PMCID: PMCPMC2685627.

8. Raote I, Malhotra V. Tunnels for Protein Export from the Endoplasmic Reticulum. Annu Rev Biochem. 2021;90:605–30. Epub 2021/01/28. doi: 10.1146/annurev-biochem-080120-022017. PubMed PMID: 33503381.

9. Liu M, Hu J, Zhang A, Dai Y, Chen W, He Y, et al. Auxilin-like protein MoSwa2 promotes effector secretion and virulence as a clathrin uncoating factor in the rice blast fungus Magnaporthe oryzae. New Phytol. 2021;230(2):720–36. Epub 2021/01/11. doi: 10.1111/nph.17181. PubMed PMID: 33423301; PubMed Central PMCID: PMCPMC8048681.

10. Qian B, Su X, Ye Z, Liu X, Liu M, Shen D, et al. MoErv29 promotes apoplastic effector secretion contributing to virulence of the rice blast fungus Magnaporthe oryzae. New Phytol. 2022;233(3):1289–302. Epub 2021/11/12. doi: 10.1111/nph.17851. PubMed PMID: 34761375; PubMed Central PMCID: PMCPMC8738142.

11. Rabouille C, Malhotra V, Nickel W. Diversity in unconventional protein secretion. J Cell Sci. 2012;125(Pt 22):5251–5. Epub 2013/02/05. doi: 10.1242/jcs.103630. PubMed PMID: 23377655.

12. Rabouille C. Pathways of Unconventional Protein Secretion. Trends Cell Biol. 2017;27(3):230–40. Epub 2016/12/19. doi: 10.1016/j.tcb.2016.11.007. PubMed PMID: 27989656.

13. Dimou E, Nickel W. Unconventional mechanisms of eukaryotic protein secretion. Curr Biol. 2018;28(8):R406–R10. Epub 2018/04/25. doi: 10.1016/j.cub.2017.11.074. PubMed PMID: 29689224.

14. Steringer JP, Bleicken S, Andreas H, Zacherl S, Laussmann M, Temmerman K, et al. Phosphatidylinositol 4,5-bisphosphate (PI(4,5)P2)-dependent oligomerization of fibroblast growth factor 2 (FGF2) triggers the formation of a lipidic membrane pore implicated in unconventional secretion. J Biol Chem. 2012;287(33):27659–69. Epub 2012/06/26. doi: 10.1074/jbc.M112.381939. PubMed PMID: 22730382; PubMed Central PMCID: PMCPMC3431657.

15. Moskes C, Burghaus PA, Wernli B, Sauder U, Durrenberger M, Kappes B. Export of Plasmodium falciparum calcium-dependent protein kinase 1 to the parasitophorous vacuole is dependent on three N-terminal membrane anchor motifs. Mol Microbiol. 2004;54(3):676–91. Epub 2004/10/20. doi: 10.1111/j.1365-2958.2004.04313.x. PubMed PMID: 15491359.

16. Kinseth MA, Anjard C, Fuller D, Guizzunti G, Loomis WF, Malhotra V. The Golgi-associated protein GRASP is required for unconventional protein secretion during development. Cell. 2007;130(3):524–34. Epub 2007/07/28. doi: 10.1016/j.cell.2007.06.029. PubMed PMID: 17655921.

17. Zhang M, Kenny SJ, Ge L, Xu K, Schekman R. Translocation of interleukin-1beta into a vesicle intermediate in autophagy-mediated secretion. Elife. 2015;4. Epub 2015/11/03. doi: 10.7554/eLife.11205. PubMed PMID: 26523392; PubMed Central PMCID: PMCPMC4728131.

18. Grieve AG, Rabouille C. Golgi bypass: skirting around the heart of classical secretion. Cold Spring Harb Perspect Biol. 2011;3(4). Epub 2011/03/29. doi: 10.1101/cshperspect.a005298. PubMed PMID: 21441587; PubMed Central PMCID: PMCPMC3062214.

19. Chen KE, Healy MD, Collins BM. Towards a molecular understanding of endosomal trafficking by Retromer and Retriever. Traffic. 2019;20(7):465–78. Epub 2019/04/18. doi: 10.1111/tra.12649. PubMed PMID: 30993794.

20. Seaman MNJ. The Retromer Complex: From Genesis to Revelations. Trends Biochem Sci. 2021;46(7):608–20. Epub 2021/02/03. doi: 10.1016/j.tibs.2020.12.009. PubMed PMID: 33526371.

21. Muhammad A, Flores I, Zhang H, Yu R, Staniszewski A, Planel E, et al. Retromer deficiency observed in Alzheimer’s disease causes hippocampal dysfunction, neurodegeneration, and Abeta accumulation. Proc Natl Acad Sci U S A. 2008;105(20):7327–32. Epub 2008/05/16. doi: 10.1073/pnas.0802545105. PubMed PMID: 18480253; PubMed Central PMCID: PMCPMC2386077.

22. Wang J, Fedoseienko A, Chen B, Burstein E, Jia D, Billadeau DD. Endosomal receptor trafficking: Retromer and beyond. Traffic. 2018;19(8):578–90. Epub 2018/04/19. doi: 10.1111/tra.12574. PubMed PMID: 29667289; PubMed Central PMCID: PMCPMC6043395.

23. Zheng W, Zhou J, He Y, Xie Q, Chen A, Zheng H, et al. Retromer Is Essential for Autophagy-Dependent Plant Infection by the Rice Blast Fungus. PLoS Genet. 2015;11(12):e1005704. Epub 2015/12/15. doi: 10.1371/journal.pgen.1005704. PubMed PMID: 26658729; PubMed Central PMCID: PMCPMC4686016.

24. Zheng H, Guo Z, Xi Y, Yuan M, Lin Y, Wu C, et al. Sorting nexin (MoVps17) is required for fungal development and plant infection by regulating endosome dynamics in the rice blast fungus. Environ Microbiol. 2017;19(10):4301–17. Epub 2017/08/25. doi: 10.1111/1462-2920.13896. PubMed PMID: 28836715.

25. Wu C, Lin Y, Zheng H, Abubakar YS, Peng M, Li J, et al. The retromer CSC subcomplex is recruited by MoYpt7 and sequentially sorted by MoVps17 for effective conidiation and pathogenicity of the rice blast fungus. Mol Plant Pathol. 2021;22(2):284–98. Epub 2020/12/23. doi: 10.1111/mpp.13029. PubMed PMID: 33350057; PubMed Central PMCID: PMCPMC7814966.

26. Rojas R, van Vlijmen T, Mardones GA, Prabhu Y, Rojas AL, Mohammed S, et al. Regulation of retromer recruitment to endosomes by sequential action of Rab5 and Rab7. J Cell Biol. 2008;183(3):513–26. Epub 2008/11/05. doi: 10.1083/jcb.200804048. PubMed PMID: 18981234; PubMed Central PMCID: PMCPMC2575791.

27. Abubakar YS, Qiu H, Fang W, Zheng H, Lu G, Zhou J, et al. FgRab5 and FgRab7 are essential for endosomes biogenesis and non-redundantly recruit the retromer complex to the endosomes in Fusarium graminearum. Stress Biology. 2021;1(1). doi: 10.1007/s44154-021-00020-3.

28. Yang C, Wang X. Lysosome biogenesis: Regulation and functions. J Cell Biol. 2021;220(6). Epub 2021/05/06. doi: 10.1083/jcb.202102001. PubMed PMID: 33950241; PubMed Central PMCID: PMCPMC8105738.

29. Borchers AC, Langemeyer L, Ungermann C. Who’s in control? Principles of Rab GTPase activation in endolysosomal membrane trafficking and beyond. J Cell Biol. 2021;220(9). Epub 2021/08/13. doi: 10.1083/jcb.202105120. PubMed PMID: 34383013; PubMed Central PMCID: PMCPMC8366711.

30. Liu XH, Chen SM, Gao HM, Ning GA, Shi HB, Wang Y, et al. The small GTPase MoYpt7 is required for membrane fusion in autophagy and pathogenicity of Magnaporthe oryzae. Environ Microbiol. 2015;17(11):4495–510. Epub 2015/05/21. doi: 10.1111/1462-2920.12903. PubMed PMID: 25991510.

31. Sweigard JA, Carroll AM, Kang S, Farrall L, Chumley FG, Valent B. Identification, cloning, and characterization of PWL2, a gene for host species specificity in the rice blast fungus. Plant Cell. 1995;7(8):1221–33. Epub 1995/08/01. doi: 10.1105/tpc.7.8.1221. PubMed PMID: 7549480; PubMed Central PMCID: PMCPMC160946.

32. Kankanala P, Czymmek K, Valent B. Roles for rice membrane dynamics and plasmodesmata during biotrophic invasion by the blast fungus. Plant Cell. 2007;19(2):706–24. Epub 2007/02/27. doi: 10.1105/tpc.106.046300. PubMed PMID: 17322409; PubMed Central PMCID: PMCPMC1867340.

33. Zarnack K, Maurer S, Kaffarnik F, Ladendorf O, Brachmann A, Kamper J, et al. Tetracycline-regulated gene expression in the pathogen Ustilago maydis. Fungal Genet Biol. 2006;43(11):727–38. Epub 2006/07/18. doi: 10.1016/j.fgb.2006.05.006. PubMed PMID: 16843015.

34. Wanka F, Cairns T, Boecker S, Berens C, Happel A, Zheng X, et al. Tet-on, or Tet-off, that is the question: Advanced conditional gene expression in Aspergillus. Fungal Genet Biol. 2016;89:72–83. Epub 2015/11/12. doi: 10.1016/j.fgb.2015.11.003. PubMed PMID: 26555930.

35. Garí E, Piedrafita L, Aldea M, Herrero E. A Set of Vectors with a Tetracycline-Regulatable Promoter System for Modulated Gene Expression inSaccharomyces cerevisiae. Yeast. 1997;13(9):837–48. doi: 10.1002/(sici)1097-0061(199707)13:9<837::Aid-yea145>3.0.Co;2-t.

36. Park YN, Morschhauser J. Tetracycline-inducible gene expression and gene deletion in Candida albicans. Eukaryot Cell. 2005;4(8):1328–42. Epub 2005/08/10. doi: 10.1128/EC.4.8.1328-1342.2005. PubMed PMID: 16087738; PubMed Central PMCID: PMCPMC1214539.

37. Qi Z, Liu M, Dong Y, Zhu Q, Li L, Li B, et al. The syntaxin protein (MoSyn8) mediates intracellular trafficking to regulate conidiogenesis and pathogenicity of rice blast fungus. New Phytol. 2016;209(4):1655–67. Epub 2015/11/03. doi: 10.1111/nph.13710. PubMed PMID: 26522477.

38. Zheng W, Lin Y, Fang W, Zhao X, Lou Y, Wang G, et al. The endosomal recycling of FgSnc1 by FgSnx41-FgSnx4 heterodimer is essential for polarized growth and pathogenicity in Fusarium graminearum. New Phytol. 2018;219(2):654–71. Epub 2018/04/21. doi: 10.1111/nph.15178. PubMed PMID: 29676464.

39. Cullen PJ, Steinberg F. To degrade or not to degrade: mechanisms and significance of endocytic recycling. Nat Rev Mol Cell Biol. 2018;19(11):679–96. Epub 2018/09/09. doi: 10.1038/s41580-018-0053-7. PubMed PMID: 30194414.

40. Perera RM, Zoncu R. The Lysosome as a Regulatory Hub. Annu Rev Cell Dev Biol. 2016;32:223–53. Epub 2016/08/09. doi: 10.1146/annurev-cellbio-111315-125125. PubMed PMID: 27501449.

41. Balderhaar HJ, Arlt H, Ostrowicz C, Brocker C, Sundermann F, Brandt R, et al. The Rab GTPase Ypt7 is linked to retromer-mediated receptor recycling and fusion at the yeast late endosome. J Cell Sci. 2010;123(Pt 23):4085–94. Epub 2010/11/11. doi: 10.1242/jcs.071977. PubMed PMID: 21062894.

42. Jahn R, Scheller RH. SNAREs--engines for membrane fusion. Nat Rev Mol Cell Biol. 2006;7(9):631–43. Epub 2006/08/17. doi: 10.1038/nrm2002. PubMed PMID: 16912714.

43. Song W, Dou X, Qi Z, Wang Q, Zhang X, Zhang H, et al. R-SNARE homolog MoSec22 is required for conidiogenesis, cell wall integrity, and pathogenesis of Magnaporthe oryzae. PLoS One. 2010;5(10):e13193. Epub 2010/10/16. doi: 10.1371/journal.pone.0013193. PubMed PMID: 20949084; PubMed Central PMCID: PMCPMC2950850.

44. Dou X, Wang Q, Qi Z, Song W, Wang W, Guo M, et al. MoVam7, a conserved SNARE involved in vacuole assembly, is required for growth, endocytosis, ROS accumulation, and pathogenesis of Magnaporthe oryzae. PLoS One. 2011;6(1):e16439. Epub 2011/02/02. doi: 10.1371/journal.pone.0016439. PubMed PMID: 21283626; PubMed Central PMCID: PMCPMC3025985.

45. Galluzzi L, Green DR. Autophagy-Independent Functions of the Autophagy Machinery. Cell. 2019;177(7):1682–99. Epub 2019/06/15. doi: 10.1016/j.cell.2019.05.026. PubMed PMID: 31199916; PubMed Central PMCID: PMCPMC7173070.

46. Chang C, Jensen LE, Hurley JH. Autophagosome biogenesis comes out of the black box. Nat Cell Biol. 2021;23(5):450–6. Epub 2021/04/28. doi: 10.1038/s41556-021-00669-y. PubMed PMID: 33903736; PubMed Central PMCID: PMCPMC8122082.

47. Raudenska M, Balvan J, Masarik M. Crosstalk between autophagy inhibitors and endosome-related secretory pathways: a challenge for autophagy-based treatment of solid cancers. Mol Cancer. 2021;20(1):140. Epub 2021/10/29. doi: 10.1186/s12943-021-01423-6. PubMed PMID: 34706732; PubMed Central PMCID: PMCPMC8549397.

48. Kershaw MJ, Talbot NJ. Genome-wide functional analysis reveals that infection-associated fungal autophagy is necessary for rice blast disease. Proc Natl Acad Sci U S A. 2009;106(37):15967–72. Epub 2009/09/01. doi: 10.1073/pnas.0901477106. PubMed PMID: 19717456; PubMed Central PMCID: PMCPMC2747227.

49. Deng YZ, Naqvi NI. Metabolic Basis of Pathogenesis and Host Adaptation in Rice Blast. Annu Rev Microbiol. 2019;73:601–19. Epub 2019/07/10. doi: 10.1146/annurev-micro-020518-115810. PubMed PMID: 31283431.

50. Ao X, Zou L, Wu Y. Regulation of autophagy by the Rab GTPase network. Cell Death Differ. 2014;21(3):348–58. Epub 2014/01/21. doi: 10.1038/cdd.2013.187. PubMed PMID: 24440914; PubMed Central PMCID: PMCPMC3921601.

51. Zhao YG, Codogno P, Zhang H. Machinery, regulation and pathophysiological implications of autophagosome maturation. Nat Rev Mol Cell Biol. 2021;22(11):733–50. Epub 2021/07/25. doi: 10.1038/s41580-021-00392-4. PubMed PMID: 34302147; PubMed Central PMCID: PMCPMC8300085.

52. Moreau K, Renna M, Rubinsztein DC. Connections between SNAREs and autophagy. Trends Biochem Sci. 2013;38(2):57–63. Epub 2013/01/12. doi: 10.1016/j.tibs.2012.11.004. PubMed PMID: 23306003.

53. Yu L, Chen Y, Tooze SA. Autophagy pathway: Cellular and molecular mechanisms. Autophagy. 2018;14(2):207–15. Epub 2017/09/22. doi: 10.1080/15548627.2017.1378838. PubMed PMID: 28933638; PubMed Central PMCID: PMCPMC5902171.

54. Hanley SE, Cooper KF. Sorting Nexins in Protein Homeostasis. Cells. 2020;10(1). Epub 2020/12/31. doi: 10.3390/cells10010017. PubMed PMID: 33374212; PubMed Central PMCID: PMCPMC7823608.

55. Deng YZ, Qu Z, He Y, Naqvi NI. Sorting nexin Snx41 is essential for conidiation and mediates glutathione-based antioxidant defense during invasive growth in Magnaporthe oryzae. Autophagy. 2012;8(7):1058–70. Epub 2012/05/09. doi: 10.4161/auto.20217. PubMed PMID: 22561104; PubMed Central PMCID: PMCPMC3429543.

56. Deng Y, Qu Z, Naqvi NI. The role of snx41-based pexophagy in magnaporthe development. PLoS One. 2013;8(11):e79128. Epub 2013/12/05. doi: 10.1371/journal.pone.0079128. PubMed PMID: 24302988; PubMed Central PMCID: PMCPMC3841179.

57. Zheng H, Chen S, Chen X, Liu S, Dang X, Yang C, et al. The Small GTPase MoSec4 Is Involved in Vegetative Development and Pathogenicity by Regulating the Extracellular Protein Secretion in Magnaporthe oryzae. Front Plant Sci. 2016;7:1458. Epub 2016/10/13. doi: 10.3389/fpls.2016.01458. PubMed PMID: 27729922; PubMed Central PMCID: PMCPMC5037964.

58. Li X, Gao C, Li L, Liu M, Yin Z, Zhang H, et al. MoEnd3 regulates appressorium formation and virulence through mediating endocytosis in rice blast fungus Magnaporthe oryzae. PLoS Pathog. 2017;13(6):e1006449. Epub 2017/06/20. doi: 10.1371/journal.ppat.1006449. PubMed PMID: 28628655; PubMed Central PMCID: PMCPMC5491321.

59. Wei YY, Liang S, Zhang YR, Lu JP, Lin FC, Liu XH. MoSec61beta, the beta subunit of Sec61, is involved in fungal development and pathogenicity, plant immunity, and ER-phagy in Magnaporthe oryzae. Virulence. 2020;11(1):1685–700. Epub 2020/11/18. doi: 10.1080/21505594.2020.1848983. PubMed PMID: 33200669; PubMed Central PMCID: PMCPMC7714445.

60. Huang L, Zhang S, Yin Z, Liu M, Li B, Zhang H, et al. MoVrp1, a putative verprolin protein, is required for asexual development and infection in the rice blast fungus Magnaporthe oryzae. Sci Rep. 2017;7:41148. Epub 2017/01/25. doi: 10.1038/srep41148. PubMed PMID: 28117435; PubMed Central PMCID: PMCPMC5259722.

61. Chen J, Zheng W, Zheng S, Zhang D, Sang W, Chen X, et al. Rac1 is required for pathogenicity and Chm1-dependent conidiogenesis in rice fungal pathogen Magnaporthe grisea. PLoS Pathog. 2008;4(11):e1000202. Epub 2008/11/15. doi: 10.1371/journal.ppat.1000202. PubMed PMID: 19008945; PubMed Central PMCID: PMCPMC2575402.

62. Khang CH, Park SY, Rho HS, Lee YH, Kang S. Filamentous Fungi (Magnaporthe grisea and Fusarium oxysporum). Methods Mol Biol. 2006;344:403–20. Epub 2006/10/13. doi: 10.1385/1-59745-131-2:403. PubMed PMID: 17033082.

63. Khang CH, Berruyer R, Giraldo MC, Kankanala P, Park SY, Czymmek K, et al. Translocation of Magnaporthe oryzae effectors into rice cells and their subsequent cell-to-cell movement. Plant Cell. 2010;22(4):1388–403. Epub 2010/05/04. doi: 10.1105/tpc.109.069666. PubMed PMID: 20435900; PubMed Central PMCID: PMCPMC2879738.

64. Park CH, Chen S, Shirsekar G, Zhou B, Khang CH, Songkumarn P, et al. The Magnaporthe oryzae effector AvrPiz-t targets the RING E3 ubiquitin ligase APIP6 to suppress pathogen-associated molecular pattern-triggered immunity in rice. Plant Cell. 2012;24(11):4748–62. Epub 2012/12/04. doi: 10.1105/tpc.112.105429. PubMed PMID: 23204406; PubMed Central PMCID: PMCPMC3531864.

