## Supplemental Tables for "Rab7/Retromer-based endolysosomal trafficking facilitates effector secretion and host invasion in rice blast"

Table S1 Putative SNARE interactors of MoVps35–GFP

| **Accession** | **Description** | **Prediction of subcellular location/trafficking** | **Biological repeat 1** | | **Biological repeat 2** | | **Biological repeat 3** | | **Biological repeat 4** | |
| --- | --- | --- | --- | --- | --- | --- | --- | --- | --- | --- |
|  |  |  | **Coverage (%)** | **Unique peptides** | **Coverage (%)** | **Unique peptides** | **Coverage (%)** | **Unique peptides** | **Coverage (%)** | **Unique peptides** |
| MGG_05089 | Vacuolar protein sorting-associated protein 35 | Endosome-golgi/PM (vesicles) | 26.50 | 26 | 65.37 | 56 | 30.85 | 19 | 52.78 | 32 |
| MGG_12614 | Hypothetical protein | Endosome-PM (vesicles) |  |  | 35.20 | 3 | 28 | 2 | 15.20 | 1 |
| MGG_00978 | Hypothetical protein | PM | 10.14 | 3 | 3.48 | 1 |  |  |  |  |
| MGG_03885 | SNARE domain-containing protein | Vacuole |  |  | 3.69 | 1 |  |  | 5.17 | 1 |
| MGG_01124 | Vesicle transport V-SNARE protein vti1 | Golgi-Vacuole |  |  |  |  | 17.17 | 2 | 9.01 | 1 |
| MGG_06521 | V-SNARE | Golgi-Vacuole |  |  | 3.99 | 1 | 6.52 | 1 | 6.52 | 1 |
| MGG_06125 | SNARE Ykt6 | Golgi-Vacuole |  |  | 4.06 | 1 |  |  | 16.75 | 2 |
| MGG_01681 | Vesicle-associated membrane protein 725 | Golgi-Vacuole |  |  | 8.86 | 2 |  |  |  |  |
| MGG_06883 | t-SNARE | Golgi |  |  | 3.92 | 1 | 12.61 | 3 | 15.13 | 3 |
| MGG_08082 | SNARE domain-containing protein | Golgi |  |  | 7.03 | 1 | 26.56 | 4 | 7.03 | 1 |
| MGG_07189 | V-SNARE | Golgi | 5.33 | 1 |  |  |  |  |  |  |
| MGG_04454 | Transporter GOS1 | ER-Golgi | 7.49 | 2 |  |  |  |  |  |  |
| MGG_12919 | Synaptobrevin | ER | 2.52 | 1 |  |  |  |  |  |  |

Table S2 Wild-type and mutant strains of fungi used in this study

| Strain | Description | Reference |
| --- | --- | --- |
| Guy11 | Wild-type (WT) | - |
| His-GFP + Pwl2-mCherry | WT strain expressing both His-GFP and Pwl2-mCherry constructs. | This study |
| GFP-MoRab7 + Pwl2-mCherry | WT strain expressing both GFP-MoRab7 and Pwl2-mCherry constructs. The GFP-MoRab7 construct was drive by the native promoter. | This study |
| GFP-MoRab7WT + Pwl2-mCherry | WT strain expressing both GFP-MoRab7 and Pwl2-mCherry constructs. The GFP-MoRab7 construct was drive by the PWL2 promoter. | This study |
| GFP-MoRab7DN(T22N) + Pwl2-mCherry | WT strain expressing both GFP-MoRab7DN(T22N) and Pwl2-mCherry constructs. The GFP-MoRab7 construct was drive by the PWL2 promoter. | This study |
| GFP-MoRab7DN(N125I) + Pwl2-mCherry | WT strain expressing both GFP-MoRab7DN(N125I) and Pwl2-mCherry constructs. The GFP-MoRab7 construct was drive by the PWL2 promoter. | This study |
| GFP-MoRab7CA(Q67L) + Pwl2-mCherry | WT strain expressing both GFP-MoRab7CA(Q67L) and Pwl2-mCherry constructs. The GFP-MoRab7 construct was drive by the PWL2 promoter. | This study |
| mCherry-MoRab7 + MoVps35-GFP | WT strain expressing both mCherry-MoRab7 and MoVps35-GFP constructs. | Wu et al. 2021 |
| GFP + MoVps35-Flag | WT strain expressing cytosolic GFP and MoVps35-Flag constructs | Wu et al. 2021 |
| GFP-MoRab7 + MoVps35-Flag | WT strain expressing both GFP-MoRab7 and MoVps35-Flag constructs. | Wu et al. 2021 |
| MoVps35-GFP + Pwl2-mCherry | WT strain expressing both MoVps35-GFP and Pwl2-mCherry constructs. | This study |
| MoVps17-GFP + Pwl2-mCherry | WT strain expressing both MoVps17-GFP and Pwl2-mCherry constructs. | This study |
| MoVps35-GFP + Bas4-mCherry | WT strain expressing both MoVps35-GFP and Bas4-mCherry constructs. | This study |
| Bas4-GFP + Pwl2-mCherry/WT | WT strain expressing both Bas4-GFP and Pwl2-mCherry constructs. | This study |
| Bas4-GFP + Pwl2-mCherry/*ΔMovps29* | *ΔMovps29* strain expressing both Bas4-GFP and Pwl2-mCherry constructs. | This study |
| Tet-Off-GFP-MoVps35/Pwl2-mCherry | Pwl2-mCherry strain expressing Tet-Off-GFP-MoVps35 construct. | This study |
| Tet-Off-GFP-MoVps35/Bas4-mCherry | Bas4-mCherry strain expressing Tet-Off-GFP-MoVps35 construct. | This study |
| GFP-MoSnc1 + MoVps35-Flag | WT strain expressing both GFP-MoSnc1 and MoVps35-Flag constructs. | This study |
| MoVps35-GFP + mCherry-MoSnc1 | WT strain expressing both MoVps35-GFP and mCherry-MoSnc1 constructs. | This study |
| WT + GFP-MoSnc1 | WT strain expressing GFP-MoSnc1 construct. | This study |
| *ΔMovps35 +* GFP-MoSnc1 | *ΔMovps35* strain expressing GFP-MoSnc1 construct. | This study |
| GFP-MoSnc1 + Pwl2-mCherry | WT strain expressing both GFP-MoSnc1 and Pwl2-mCherry constructs. | This study |
| GFP-MoSnc1 + Bas4-mCherry | WT strain expressing both GFP-MoSnc1 and Bas4-mCherry constructs. | This study |
| Bas4-GFP + Pwl2-mCherry/*ΔMosnc1* | *ΔMosnc1* strain expressing both Bas4-GFP and Pwl2-mCherry constructs. | This study |
| *ΔMosnc1* | *MoSNC1* deletion mutant of WT. | This study |
| *ΔMosnc1-C* | *ΔMosnc1* strain expressing GFP-MoSnc1 construct | This study |
| WT + Bas4-mCherry | WT strain expressing Bas4-mCherry construct. | This study |
| GFP-MoRab7DN(T22N) + Bas4-mCherry | WT strain expressing both GFP-MoRab7DN(T22N) and Bas4-mCherry constructs. | This study |
| GFP-MoRab7DN(N125I) + Bas4-mCherry | WT strain expressing both GFP-MoRab7DN(N125I) and Bas4-mCherry constructs. | This study |
| MoVps35-nYFP + Pwl2-cYFP | WT strain expressing both MoVps35-nYFP and Pwl2-cYFP constructs. | This study |
| MoVps35-GFP | *ΔMovps35* strain expressing MoVps35-GFP construct. | Zheng et al., 2015 |

Table S3 Plasmids used in this study.

| Plasmid | Description |
| --- | --- |
| Pwl2-mCherry | *PWL2* (MGG_04301) promoter and entire coding sequence with a C-terminal translational fusion of the mCherry reporter gene. The *PWL2* gene fragment cloned into KpnI-HindIII restriction sites of pKNT-mCherry (Geneticin^R^, Ampicillin^R^). |
| His-GFP | *CCG1* promoter and *N. crassa* histone H1-GFP cassette with *A. nidulans* TrpC terminator. Published in Yang et al., 2014. |
| GFP-MoRab7 | *MoRAB7* (MGG_08144) promoter and entire coding sequence with a N-terminal translational fusion of the GFP reporter gene. Published in Wu et al., 2021. |
| mCherry-MoRab7 | *MoRAB7* (MGG_08144) promoter and entire coding sequence with a N-terminal translational fusion of the mCherry reporter gene. Published in Wu et al., 2021. |
| GFP-MoRab7WT | The GFP-MoRab7WT fragment was amplified from above plasmid (GFP-MoRab7) and subsequently fused with the PWL2 promoter. The final PWL2::GFP::MoRab7WT fragment cloned into KpnI-HindIII restriction sites of pKNT (Geneticin^R^, Ampicillin^R^). |
| GFP-MoRab7DN(T22N) | The GFP-MoRab7DN(T22N) fragment was amplified from previous plasmid that has been published by Wu. et al., 2021.  The generated fragment was fused with the PWL2 promoter. The final PWL2::GFP::MoRab7DN(T22N) fragment cloned into KpnI-HindIII restriction sites of pKNT (Geneticin^R^, Ampicillin^R^). |
| GFP-MoRab7DN(N125I) | The GFP-MoRab7DN(N125I) fragment was amplified from previous plasmid that has been published by Wu. et al., 2021.  The generated fragment was fused with the PWL2 promoter. The final PWL2::GFP::MoRab7DN(N125I) fragment cloned into KpnI-HindIII restriction sites of pKNT (Geneticin^R^, Ampicillin^R^). |
| GFP-MoRab7CA(Q67L) | The GFP-MoRab7CA(Q67L) fragment was amplified from previous plasmid that has been published by Wu. et al., 2021.  The generated fragment was fused with the PWL2 promoter. The final PWL2::GFP::MoRab7CA(Q67L) fragment cloned into KpnI-HindIII restriction sites of pKNT (Geneticin^R^, Ampicillin^R^). |
| MoVps35-GFP | *MoVPS35* (MGG_05089) promoter and entire coding sequence with a C-terminal translational fusion of the GFP reporter gene. Published in Zheng et al., 2015. |
| MoVps35-Flag | *MoVPS35* (MGG_05089) promoter and entire coding sequence with a C-terminal translational fusion of the Flag tag. Published in Wu et al., 2021. |
| MoVps17-GFP | *MoVPS17* (MGG_01434) promoter and entire coding sequence with a C-terminal translational fusion of the GFP reporter gene. Published in Zheng et al., 2017. |
| Tet-Off-GFP-MoVps35 | See Materials and Methods (Tet-Off gene expression system) |
| Bas4-GFP | *BAS4* (MGG_10914) promoter and entire coding sequence with a C-terminal translational fusion of the GFP reporter gene. The BAS4 gene fragment cloned into KpnI-HindIII restriction sites of pKNT-GFP (Geneticin^R^, Ampicillin^R^). |
| Bas4-mCherry | *BAS4* (MGG_10914) promoter and entire coding sequence with a C-terminal translational fusion of the mCherry reporter gene. The BAS4 gene fragment cloned into KpnI-HindIII restriction sites of pKNT-mCherry (Geneticin^R^, Ampicillin^R^). This construction (Bas4-mCherry) also been cloned into the pFGL1010 vector (Sulfonylurea^R^, Kanamycin^R^). |
| GFP-MoSnc1 | *MoSNC1* (MGG_12614) promoter and entire coding sequence with a N-terminal translational fusion of the GFP reporter gene (Geneticin^R^, Ampicillin^R^). |
| mCherry-MoSnc1 | *MoSNC1* (MGG_12614) promoter and entire coding sequence with a N-terminal translational fusion of the GFP reporter gene (Geneticin^R^, Ampicillin^R^). |
| MoVps35-nYFP | *MoVPS35* (MGG_05089) promoter and entire coding sequence with a C-terminal translational fusion of the nYFP reporter gene. A 4.7-kb *MoVPS35* gene fragment cloned into KpnI-HindIII restriction sites of pKNT-NYFP (Geneticin^R^, Ampicillin^R^). |
| Pwl2-cYFP | *PWL2* (MGG_04301) promoter and entire coding sequence with a C-terminal translational fusion of the cYFP reporter gene. A 1.3-kb *PWL2* gene fragment cloned into KpnI-HindIII restriction sites of pCX62-CYFP (Hygromycin^R^, Ampicillin^R^). |

Table S4 PCR primers used in this study

| Primers | Sequence(5’-3’) | Application |
| --- | --- | --- |
| PWL2-PF | agggaacaaaagctgggtaccGCGTCAGTGAACAAACC | Construction of Pwl2-mCherry |
| PWL2-PR | gcccttgctcaccataagcttCATAATATTGCAGCCCTC |  |
| P-PWL2-F | agggaacaaaagctgggtaccGCGTCAGTGAACAAACCTG | Construction of PWL2::GFP::MoRab7WT, PWL2::GFP::MoRab7DN(T22N), PWL2::GFP::MoRab7DN(N125I) and PWL2::GFP::MoRab7CA(Q67L) vector |
| P-PWL2-R | AACAGCTCCTCGCCCTTGCTCACCATTTTGAAAGTTTTTAATTTTAAAAAG |  |
| G-Rab7DN-F | ATGGTGAGCAAGGGCGAG |  |
| G-Rab7DN-R | gggctgcaggcatgcaagcttTTAGCAGGCGCATCCATC |  |
| BAS4-PF | agggaacaaaagctgggtaccGGTAGCTTCTACGGATGCGT | Construction of Bas4-GFP or Bas4-mCherry vector (Geneticin^R^) |
| BAS4-PR | gcccttgctcaccataagcttAGCAGGGGGGATAGACGA |  |
| BAS4-mC-CF | catgatgatgctcgagaattcGGTAGCTTCTACGGATGCG | Construction of Bas4-mCherry vector Sulfonylurea^R^) |
| BAS4-mC-CR | gactctagaactagtggatccCTACTTGTACAGCTCGTCCATG |  |
| MoVps35-nYFP-F | agggaacaaaagctgggtaccTGCCCAATGTTACAAGGTGC | Construction of MoVps35-nYFP vector |
| MoVps35-nYFP-R | cgtggcgatggagcgaagcttCTTGGGATCCAACACAATTCC |  |
| Pwl2-cYFP-F | agggaacaaaagctgggtaccGCGTCAGTGAACAAACCTG | Construction of Pwl2-cYFP vector |
| Pwl2-cYFP-R | cttgcaggccgggcgaagcttCATAATATTGCAGCCCTCTTC |  |
| MoVps35Tet-AF | attattatggagaaactcgagTGGCAGATGCTTCTCACTCAG | Construction of Tet-Off-GFP-MoVps35 vector |
| MoVps35Tet-AR | atccaggcgggccatgaattcTGGTTCTGGCTGGCTGAACT |  |
| MoVps35Tet-BF | agttctagagtcgacctgcagATGGCGTCGGTCCCAGCTC |  |
| MoVps35Tet-BR | acgacggccagtgccaagcttACTCACGCTCGGCATAATCG |  |
| GFP-F | ATGGTGAGCAAGGGCGAG | Identification of Tet-Off-GFP-MoVps35 transformants |
| MoVps35-OR | TCTTTACTGGCTCCTGTTGCTC |  |
| MoVps35UA-F | GGCTTCCCAGCAACCTTTC | Identification of Tet-Off-GFP-MoVps35 transformants |
| tTA-R | CCTCGTTCAGCAGCTCCAA |  |
| MGG_12614AF | GAGATTATGCTCCTACCGC | Gene deletion of *MoSNC1* (MGG_12614) |
| MGG_12614AR | TTGACCTCCACTAGCTCCAGCCAAGCCGATGGATGGGGATGAAAA |  |
| MGG_12614BF | GAATAGAGTAGATGCCGACCGCGGGTTATGAATCCCTCGGTTTGT |  |
| MGG_12614BR | CTGTTGTCTTGCGTGCTG |  |
| MGG_12614OF | AAGACGCTCCCTACGACC | Identification of *ΔMosnc1* |
| MGG_12614OR | GCCCTTGAAGTGGAAAAC |  |
| MGG_12614UA | ACTTACCCCTGAGTGCTTC | Identification of *ΔMosnc1* |
| H853 | GACAGACGTCGCGGTGAGTT |  |
| MGG_12614PF1 | gtcgacggtatcgataagcttGGCAACATCCATAGGGCATC | Construction of GFP-MoSnc1 vector |
| MGG_12614PR1 | GAACAGCTCCTCGCCCTTGCTCACCATGTTTGCGGTTGCGGCTCTTT |  |
| eGFP-OF | ATGGTGAGCAAGGGCGAG |  |
| eGFP-OR | CTTGTACAGCTCGTCCAT |  |
| MGG_12614ORF-F1 | ACTCACGGCATGGACGAGCTGTACAAGATGCCCGAAGACGCTCCCTA |  |
| MGG_12614ORF-R1 | tcagtaacgttaagtggatccAGGGTTGCTCCGTTCCGTC |  |
| MGG_12614PF2 | catgatgatgctcgagaattcGGCAACATCCATAGGGCATC | Construction of mCherry-MoSnc1 vector |
| MGG_12614PR2 | GTTATCCTCCTCGCCCTTGCTCACCATGTTTGCGGTTGCGGCTCTTT |  |
| mCherry-OF | ATGGTGAGCAAGGGCGAG |  |
| mCherry-OR | CTTGTACAGCTCGTCCATGC |  |
| MGG_12614ORF-F2 | ACCGGCGGCATGGACGAGCTGTACAAGATGCCCGAAGACGCTCCCTA |  |
| MGG_12614ORF-R2 | gactctagaactagtggatccAGGGTTGCTCCGTTCCGTC |  |
